# Ubiquitin proteomics uncovers RNA polymerase I as a critical target of the Nse1 RING domain

**DOI:** 10.1101/2021.12.11.472054

**Authors:** Eva Ibars, Joan Codina-Fabra, Gemma Bellí, Celia Casas, Marc Tarrés, Roger Solé-Soler, Neus P. Lorite, Pilar Ximénez-Embún, Javier Muñoz, Neus Colomina, Jordi Torres-Rosell

**Affiliations:** Departament de Ciencies Mediques Basiques, Institut de Recerca Biomedica de Lleida, Universitat de Lleida, 25198 Lleida, Spain; Proteomics Unit. Spanish National Cancer Research Center (CNIO), Madrid, Spain; ProteoRed-ISCIII. Madrid, Spain; Biocruces Bizkaia Health Research Institute, Barakaldo, Spain; Ikerbasque, Basque Foundation for Science. Bilbao, Spain

## Abstract

Ubiquitination controls numerous cellular processes, and its deregulation is associated to many pathologies. The Nse1 subunit in the Smc5/6 complex contains a RING domain with ubiquitin E3 ligase activity and essential functions in genome integrity. However, Nse1-dependent ubiquitin targets remain largely unknown. Here, we use label-free quantitative proteomics to analyse the nuclear ubiquitinome of *nse1-C274A* RING mutant cells. Our results show that Nse1 impacts on the ubiquitination of several proteins involved in DNA damage tolerance, ribosome biogenesis and metabolism that, importantly, extend beyond canonical functions of the Smc5/6 complex in chromosome disjunction. In addition, our analysis uncovers an unexpected connection between Nse1 and RNA polymerase I (RNAP I) ubiquitination. Specifically, Nse1 and the Smc5/6 complex promote the ubiquitination of K408 and K410 in the clamp domain of Rpa190, a modification that induces its degradation in response to blocks in transcriptional elongation. We propose that this mechanism contributes to Smc5/6-dependent segregation of the rDNA array, the locus transcribed by RNAP I.

## INTRODUCTION

Post-translational modification of proteins is one of the most frequently used mechanisms to control protein function. These modifications can trigger conformational changes in the substrate, new interactions, or change the subcellular localization and stability of proteins. Ubiquitination can modify proteins through the formation of an isopeptide bond between the C-terminal glycine residue of ubiquitin (Ub) and the ε-amino group of a lysine in the target protein ^1^. Ub itself can also be targeted for ubiquitination. To this end, it uses one of its seven lysines as acceptor sites, giving rise to a wide diversity of ubiquitin chains and possible signaling modes^1^. Depending on the type of Ub chain decoration of the substrate, ubiquitination may signal for proteasomal degradation, DNA repair, protein trafficking or extraction from a specific compartment or protein complex. Ubiquitination depends on an enzymatic cascade, involving a Ub activating enzyme (E1), a Ub conjugating enzyme (E2) and a Ub ligase enzyme (E3). In budding yeast, one E1, eleven E2s and a large family of E3 enzymes catalyze the transfer of Ub molecules to specific substrates ^2^. Protein ubiquitination is balanced by ubiquitin-specific proteases (UBPs), which cleave the isopeptide bond between Ub and the target thus reversing the modification ^3^. Although ubiquitination is regulated at multiple levels, E3 enzymes are strategic players, as they provide specificity to the ubiquitination reaction. Most E3s catalyze the transfer of ubiquitin by bridging the interaction between a Ub-charged E2 enzyme and the substrate. The most abundant type of E3 ligases contain RING domains, which interact with the E2 and reorient the E2-Ub thioester bond to favor the discharge of Ub onto the substrate ^4^. Proper folding of RING domains requires the presence of eight conserved cysteine or histidine residues coordinating two zinc atoms in a cross-brace conformation ^5^. This folding is essential for their E3 activity, and mutation of these residues severely affects ubiquitination of their targets.

Ubiquitination plays crucial roles in DNA repair, by enabling extensive signaling in response to DNA damage and replicative stress ^6, 7^. The maintenance of genome integrity also requires the action of the Structural Maintenance of Chromosome (SMC) complexes, present in all domains of life. SMC complexes use their ATPase activity to form loops and organize genomes during chromosome replication, segregation and DNA repair. Particularly, the Smc5/6 complex enables chromosome disjunction by promoting removal of DNA-mediated connections between sister chromatids, which mainly arise during DNA replication and recombinational repair^8, 9^.

The Smc5/6 complex is assembled from eight subunits: Smc5 and Smc6, which constitute the core of the complex, and six non-SMC subunits (Nse proteins), Nse1 to Nse6. Unlike other SMC complexes, Smc5/6 contains two RING domains, located in the C-terminal part of the Nse1 and Nse2 subunits ^10^. The RING domain in Nse2 has SUMO ligase activity, while the RING domain in Nse1 has Ub ligase activity ^11–15^. Nse1 interacts with Nse3, a member of the highly conserved MAGE (melanoma-associated antigen) family of proteins through its N-terminal domain. The central part of Nse1 associates with Nse4, a member of the kleisin family of SMC subunits ^16^, to form a subcomplex that then binds to the ATPase domain of Smc5 and the neck region of Smc6. The C-terminal RING domain in Nse1 homologues (NH-RING) is characterized by the presence of a C4HC3 configuration of zinc-coordinating residues, closely related to the PHD (Plant Homology) domain and different from the conventional C3HC4 consensus found in other E3 RING domains ^17^. Nse1 proteins show other features of Ub E3 ligases, including the capacity to form Ub chains in vitro ^14, 15^. In vitro, Nse1 ubiquitinates Nse3 and Nse4, and its E3 ligase activity is enhanced by Nse3 ^14, 18, 19^. In human cells, Nse1 targets Mms19 in an Smc5/6-independent manner for its ubiquitination and degradation, with relevant implications in DNA polymerase function and genome stability ^20^. However, and differently to Nse2, no other in vivo targets for Nse1 have been identified until now. Although the *NSE1* RING domain is not essential for viability, it contributes to the DNA repair functions of the Smc5/6 complex, and *nse1* mutant cells carrying mutations in zinc- coordinating residues show an aberrant morphology and severe growth defects, particularly under DNA damaging conditions ^17, 21, 22^.

To fill the gap in our current knowledge of Nse1-dependent Ub targets, we have compared the nuclear ubiquitinome of wild type yeast cells and *nse1* mutants bearing a C274A mutation in the RING domain ^21^. Our analysis reveals that Nse1 affects the ubiquitination status of several proteins involved in ribosome biogenesis and metabolism, extending beyond canonical functions of the Smc5/6 complex in chromosome segregation. Moreover, Nse1 promotes the ubiquitination Rpa190, the largest subunit in RNA Polymerase I (RNAP I), at K408 and K410, a modification that promotes Rpa190 destruction. In addition, we show that this modification is triggered by blocks in RNAP I elongation, including those inflicted by UV-light. Overall, we propose that Rpa190 ubiquitination clears stalled RNAP I complexes from rDNA chromatin to facilitate other chromosomal transactions in this locus.

## RESULTS

### A proteomic screen to identify Nse1-sensitive ubiquitin sites

To identify Nse1-sensitive ubiquitin sites, we profiled the ubiquitinome of wild type and *nse1-C274A* mutant cells using diGly proteomics ^23^. The latter are affected in a conserved zinc-coordinating cysteine residue in the RING domain ^21^. As *nse1- C274A* mutant cells are sensitive to DNA damage, and to maximize differences with their wild type counterpart, exponentially growing cultures were treated with 0.02% MMS for 1.5 hours. In addition, we fractionated whole cell extracts into soluble and nuclear fractions to reduce the complexity of the sample. Protein extracts from the nuclear fraction were trypsinized and peptides containing the signature di-glycine remnant from ubiquitin were enriched by immunopurification. Finally, ubiquitinated peptides from two biological and two technical replicates were quantified by label-free mass-spectrometry (Figure 1A)^24^. We used the flowthrough from the immunoprecipitation to quantify the total protein content in the nuclear extract, allowing the identification of 3657 proteins (Supplementary Data 1). 62 proteins were significantly less expressed (q-value<0.05) and 207 proteins were upregulated in *nse1* samples (Supplementary Figure 1B). Proteins showing higher expression in *nse1* mutant cells were enriched in functions related to purine ribonucleotide metabolism and cellular respiration (FDR 4.32E-2 and 3.90E-2). One of the proteins showing the highest expression in *nse1* mutant cells was Leu2, the auxotrophic marker used for selection of the *nse1-C274A* allele, and an internal control for the total protein quantitation (Supplementary Figure 1B). Proteins less expressed in the *nse1* mutant cells were enriched in functions related to amino acid metabolism (FDRs 2.84E-06 to 3.60E-02). In addition, three proteins encoded by 2μ plasmids (Rep1, Rep2 and Raf1) were downregulated in the *nse1-C274A* mutant, suggesting that the mutation might restrict the maintenance of episomal plasmids. Interestingly, a recent screen for mutants defective in maintenance of 2μ plasmids identified a variant in the Nse2 subunit of the Smc5/6 complex ^25^.

**Figure 1.**
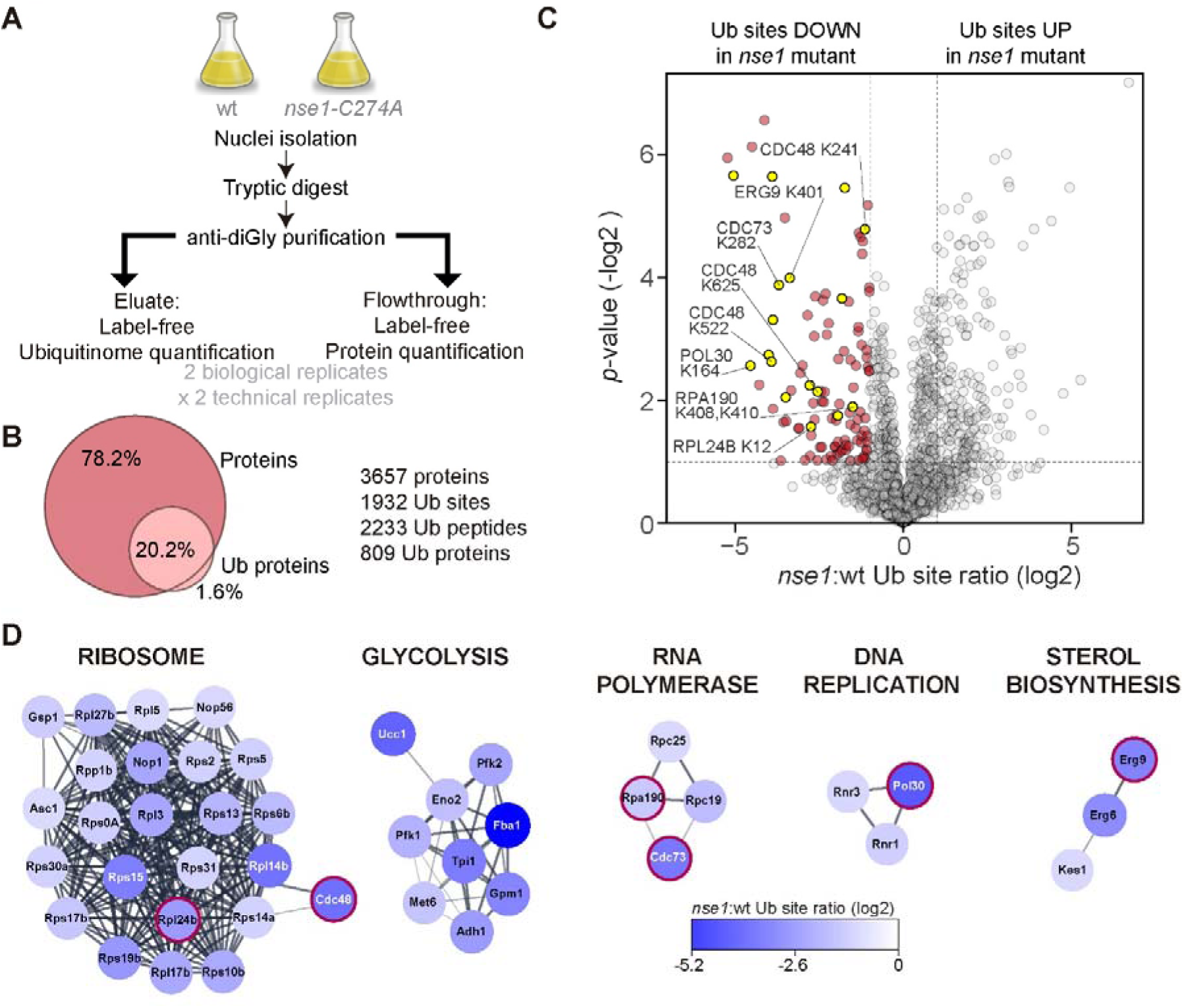
A proteomic screen to identify Nse1-sensitive ubiquitin sites. **A.** Experimental workflow for label-free quantification of ubiquitin sites and total protein levels in nuclear fractions form wild type and *nse1-C274A* mutant cells. Two biological and two technical replicates were used for each strain. **B**. Venn diagram of proteins identified in nuclear extract (Proteins) and in anti-diGly purification (Ub proteins) and summary of number of proteins, peptides and sites identified. **C.** Volcano plot showing the differentially ubiquitinated sites (log_2_ ratios) between wild type and *nse1- C274A* mutant cells on the x-axis and their statistical significance (−log_10_(*p*-value)) on the y-axis; significantly affected sites are shown above the dotted line; the ubiquitin sites downregulated in *nse1-C274A* cells are shown in red on the left side; the position of specific ubiquitination sites is shown in yellow in the graph; yellow dots without label correspond to Ub peptides from the Yhb1 protein. **D.** Cytoscape Markov cluster analysis of ubiquitin sites significantly downregulated in *nse1- C274A* cells. Note the presence of a cluster of (mainly) ribosomal proteins, proteins involved in glycolytic functions, RNA polymerase subunits, DNA replication factors and ergosterol biosynthesis enzymes. Red circles highlight targets analyzed through ubiquitin pull downs in this study.

809 of the proteins in the flowthrough (around 20%) were also found after di-Gly immunoprecipitation (Figure 1B). In total, we identified 1932 ubiquitinated sites. The nuclear enrichment step used in our protocol most probably accounts for the lower number of ubiquitinated peptides identified, relative to other studies ^26, 27^ (Supplementary Figure 1C). As shown in Figure 1C, 96 diGly-sites were significantly (q-value<0.1) less ubiquitinated in *nse1-C274A* mutant cells, representing potential Nse1-Smc5/6-dependent ubiquitin targets. Gene ontology analysis revealed that peptides with lower levels of ubiquitination in *nse1-C274A* cells were enriched in categories related to glycolysis, sterol synthesis, proteasome-mediated destruction, ribosomal proteins and RNA polymerases (Supplementary Figure 1D). Markov cluster (MCL) analysis of Nse1-dependent high confidence hits revealed the presence of several subclusters, with nodes of ribosomal, glycolysis, RNA polymerase, and sterol biosynthesis proteins (Figure 1D). In addition, we identified one subcluster, containing DNA replication factors, which included Pol30 and ribonucleotide reductase subunits.

Next, we selected seven potential Nse1-dependent ubiquitin targets from different subclusters and subjected them to further validation: Rpl24b, a 60S ribosomal subunit protein; Cdc73, a component of the Paf1 transcription elongation complex; Yhb1, a yeast flavohemoglobin involved in oxidative stress responses; Erg9, also known as squalene synthase, an enzyme that participates early in the sterol biosynthesis pathway; Cdc48, a ubiquitin-selective segregase able to extract proteins from larger cellular structures; and Pol30, a replication factor also known as sliding clamp or proliferating cell nuclear antigen (PCNA). In addition, we also confirmed Nse1-dependent ubiquitination of Rpa190 (see below). Ubiquitination of Pol30 at K164 in response to DNA damage has been extensively documented and plays important roles in genomic integrity ^28^. In contrast, the role of Rpl24b, Cdc73, Yhb1, Erg9 or Cdc48 ubiquitination is unknown. Each target was tagged with a C-terminal 6xHA epitope at its endogenous location except Pol30, which was analyzed with specific antibodies. Besides, we introduced a multicopy plasmid expressing a 7xHis tagged version of ubiquitin from the *CUP1* promoter. Protein extracts from exponentially growing cells treated or not with 0,02% MMS for 1,5 hours were prepared under denaturing conditions and ubiquitinated proteins were pulled down using NiNTA beads and analyzed by western blot. As shown in Figure 2, ubiquitination of targets was either not detectable or severely impaired in *nse1- C274A* cells, relative to wild type cells, confirming our proteomic data. For Rpl24b, Cdc73 and Erg9, MMS-induced DNA damage had little effect on ubiquitination levels (Figure 2A-C). In contrast, treatment of cells with MMS upregulated the ubiquitination of Yhb1, Cdc48 and, as previously known, Pol30 (Figure 2D-E). Thus, Nse1- dependent ubiquitin targets do not respond similarly to DNA alkylation damage.

**Figure 2.**
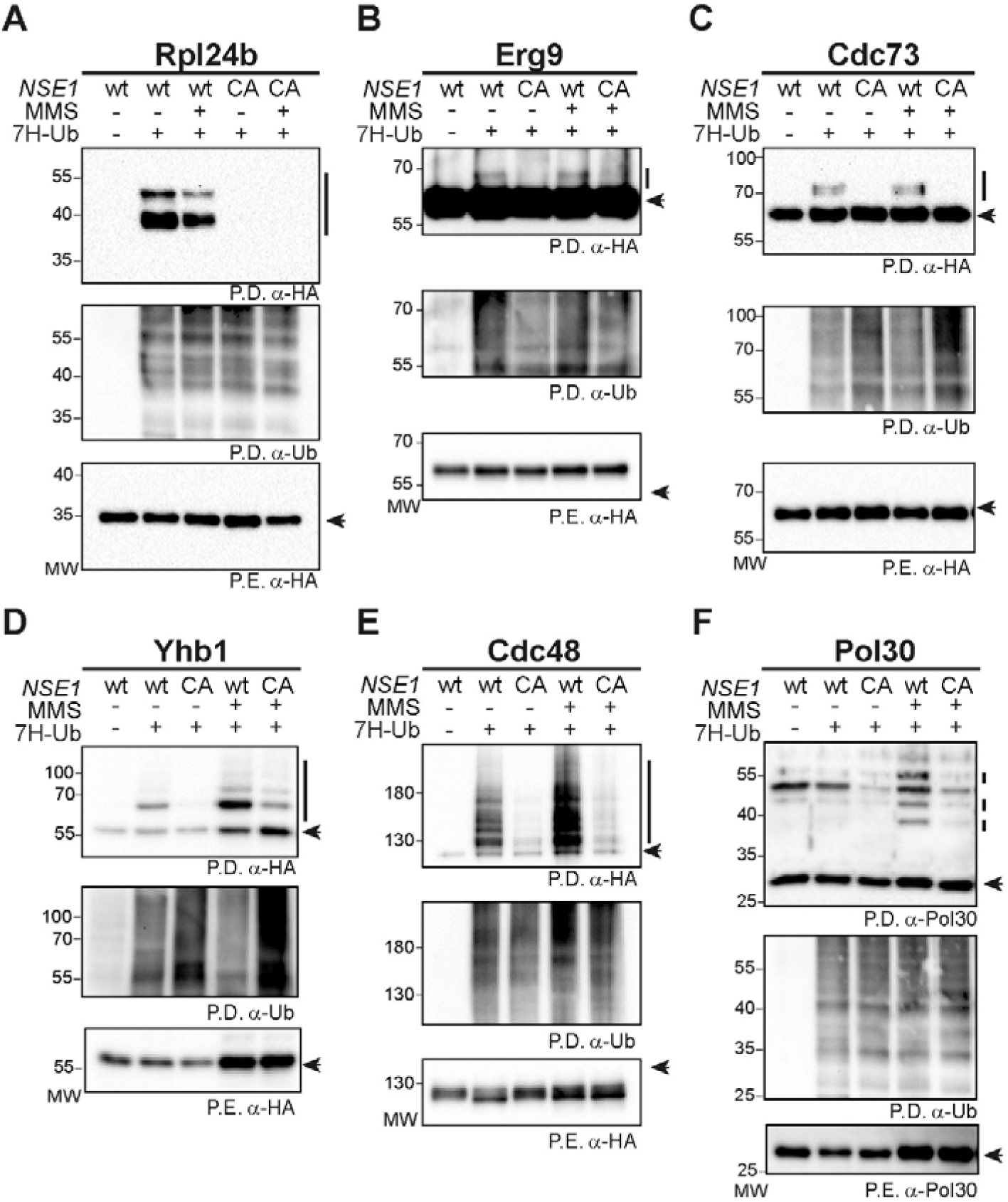
Validation of Nse1-dependent ubiquitin targets. Exponentially growing cultures of wild type (wt) or *nse1-C274A* mutant (CA) cells, expressing a C-terminal 6xHA tagged version of the indicated proteins from its endogenous location, and expressing (+) or not (-) a 7xHis tagged version of ubiquitin (7H-Ub) from the *CUP1* promoter, were collected; where indicated, cultures were treated with MMS 0.02% for 90 min before collection; ubiquitinated species from protein extracts (P.E.) were analyzed by pull down (P.D.) under denaturing conditions. 6HA-tagged proteins were detected with anti-HA antibodies; Pol30 was detected with a rabbit polyclonal antibody.

Overall, the altered ubiquitin site usage in *nse1-C274A* mutant cells indicates that Nse1 participates in various cellular processes that, importantly, extend beyond canonical Smc5/6 functions in chromosome replication, segregation and repair.

### Ubiquitination of lysine 408 and 410 modulates Rpa190 function

Another group of Nse1-dependent ubiquitin targets revealed in our data comprises RNA polymerase subunits. This cluster includes one subunit of RNAP I (Rpa190), a subunit shared by RNAP I and III complexes (Rpc19) and an RNAP III specific subunit (Rpc25). The ubiquitin site showing the greatest difference between wild type and *nse1* mutants is K122 in Rpc19, followed by the double ubiquitination of K408 and K410 in Rpa190 (Supplementary Figure 2). Of note, Smc5/6, RNAP I and RNAP III localize and operate in the rDNA array ^29, 30^, and inactivation of RNAP I alleviates the rDNA non-disjunction defects of Smc5/6 thermosensitive mutants ^8^. Therefore, we centered our interest in Rpa190, the largest subunit in RNAP I.

The K408 and K410 residues are located on top of the clamp module in the RNAP I complex (Figure 3A), a region that participates in the transition between the open and closed conformation of the polymerase ^31, 32^. Of note, sequence and structure alignment of yeast Rpa190 and human Rpa194 indicate that K408 is conserved in evolution and is occupied by K350 in the human RNAP I complex (Supplementary Figure 3A and 3B) ^33^. Rpa190 has a prominent ubiquitination band, with an electrophoretic mobility equivalent to a mass increase of around 29 KDa, which might reflect the addition of 3 ubiquitin molecules (Figure 3A); besides, there are two extra fainter bands with a mass increase comparable to the addition of 4 and 6 ubiquitin molecules, respectively. To study the effect of ubiquitination at K408/K410, we mutated both residues to arginine, generating an *rpa190-K408R,K410R* (hereafter referred to as *rpa190-KR*) allele. This mutant was first expressed from a centromeric plasmid in an *RPA190/rpa190Δ* diploid strain. Tetrad analysis of the progeny showed that *rpa190-KR* mutants are viable and not affected in growth. Next, we integrated the *rpa190-KR* allele and studied the ubiquitination status of the mutant protein using ubiquitin pull downs. As shown in Figure 3B, Rpa190 ubiquitination is severely impaired in *rpa190-KR* cells, with almost no ubiquitination forms detected in the western blot, indicating that ubiquitin mainly targets K408 and K410 in Rpa190.

**Figure 3.**
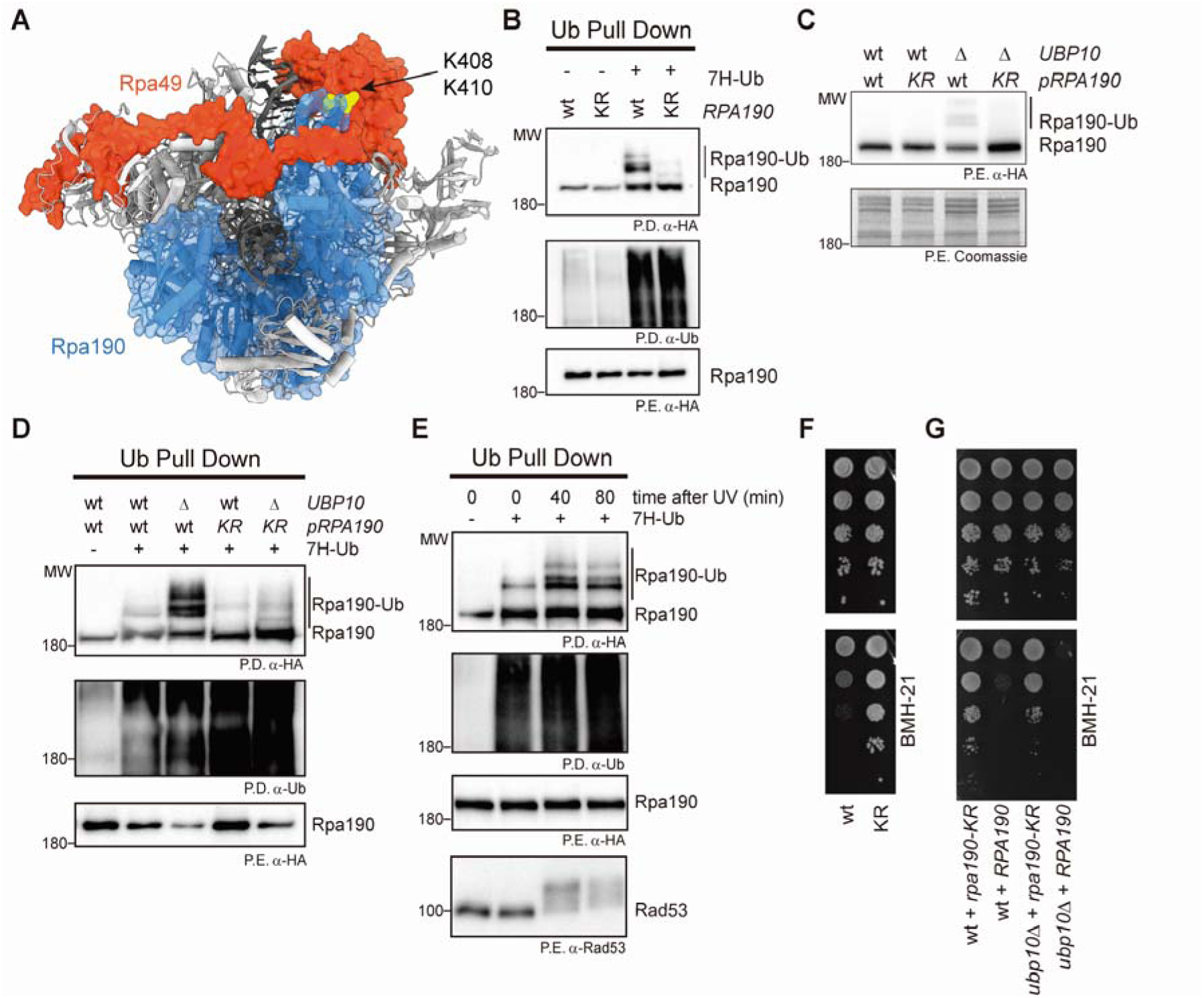
Ubiquitination of lysines 408 and 410, at the tip of the clamp domain, modulates Rpa190 protein levels and function. **A.** Structure (PDB 6H68) of an elongating Pol I complex ^38^. Rpa190 is shown in blue, Rpa49 in red; K408 and K410 residues in the clamp domain are highlighted in yellow. rDNA and rRNA are shown in dark grey. **B.** Exponentially growing cultures of the indicated strains, expressing (+) or not (-) a 7xHis tagged version of ubiquitin (7H-Ub) from the *CUP1* promoter, were collected; ubiquitinated species from protein extracts (P.E.) were enriched by pull down (P.D.) of ubiquitin under denaturing conditions. **C.** Western blot analysis of wild type (wt) and *ubp10Δ* (*Δ*) mutant cells, expressing the indicated *RPA190* alleles tagged with 6xHA; wt=wild type, KR=*rpa190-KR*; bottom panel, coomassie staining of the membrane**. D.** Pull down analysis of Rpa190-6xHA ubiquitination in wild type (wt) and *ubp10Δ* (*Δ*) mutant cells expressing the indicated *RPA190* alleles; wt=wild type, KR=*rpa190-KR*; 7H-Ub=7xHis tagged ubiquitin. **E.** Ubiquitin pull down analysis in samples from an exponentially growing culture of wild type cells expressing Rpa190-6xHA, untreated or at the indicated times after UV-irradiation (300 J/m^2^). UV-dependent activation of the DNA damage checkpoint was detected by the appearance of lower mobility Rad53 phosphorylated forms. **F**. Growth test analysis of wild type (wt) and *rpa190-KR* (KR); 10-fold serial dilutions of the liquid cultures were spotted on YPD (up) or YPD with BMH-21 15 μM (bottom). **G.** Wild type (wt) and *ubp10Δ* cells transformed with a centromeric vector expressing the indicated *RPA190* alleles were grown overnight in selective media and spotted on YPD (up) or YPD with 15 μM BMH-21 (bottom); pictures were taken after 48 hours. In B, C, D and E, 6xHA-tagged Rpa190 was detected with anti-HA antibodies; ubiquitinated forms are indicated with a vertical bar.

Rpa190 is deubiquitinated by the Ubp10 ubiquitin-specific peptidase, preventing its proteasomal degradation ^34^. In accordance, Rpa190 protein levels decreased in *ubp10Δ* cells (Figure 3C). Remarkably, the *rpa190-KR* allele restored Rpa190 protein levels in *ubp10Δ* mutant cells (Figure 3C). To test if Rpa190 protein and ubiquitination levels are inversely correlated, we performed ubiquitin pull downs in wild type and *ubp10Δ* cells expressing *RPA190* or the *rpa190-KR* alleles from a centromeric plasmid. As previously described ^34^, deletion of *UBP10* led to higher levels of Rpa190 ubiquitination (Figure 3D), an effect that could be also directly detected in protein extracts (Figure 3C). In contrast, ubiquitination was impaired in the Rpa190-KR mutant protein, a situation that was more evident in *ubp10Δ* cells (Figure 3D). Overall, these results demonstrate that ubiquitination at lysines K408 and K410 targets Rpa190 for destruction, a process that is normally counteracted by the Ubp10 deubiquitinase.

Next, we explored situations that might promote Rpa190 ubiquitination. Rpa190 ubiquitination did not change in cells with high (190) or low (25) rDNA copy number ^35^, suggesting that the relative number of actively-transcribed copies of rDNA (higher in the 25 copy number strain) does not affect Rpa190 ubiquitination (Supplementary Figure 4A). rDNA copy number in these strains is maintained through deletion of the *FOB1* gene, which normally prevents collisions between replication and RNAP I-dependent transcription, further suggesting that head-on encounters between the two machineries are also not responsible for Rpa190 ubiquitination. Rpa190 ubiquitination did not change at different stages in the cell cycle, monitored by arresting cells in G1 with alpha factor, in S phase with hydroxyurea or in G2/M with nocodazole (Supplementary Figure 4B). On the other hand, MMS-induced alkylation DNA damage slightly increased Rpa190 ubiquitination (Supplementary Figure 4B), while irradiation of cells with UV light promoted the accumulation of ubiquitinated Rpa190 species (Figure 3E and Supplementary Figure 4B), suggesting that Rpa190 is modified in response to bulky lesions on DNA, which are most probably encountered by RNAP I during transcriptional elongation. However, UV-induced ubiquitination was not accompanied by a reduction in Rpa190 protein levels (Figure 3E). Thus, either UV-dependent Rpa190 ubiquitination does not signal for its proteasomal degradation, or only a small pool of stalled RNAP I complexes are targeted by ubiquitin for destruction in response to UV damage. In addition, *rpa190- KR* mutant cells were not sensitive to UV irradiation or alkylation DNA damage (Supplementary Figure 4C), suggesting that Rpa190 ubiquitination might act redundantly with other mechanisms for repair of DNA lesions.

To further explore the effects of a transcriptional elongation block on Rpa190 ubiquitination, we analyzed the sensitivity of *rpa190-KR* cells to BMH-21, a specific inhibitor of RNAP I ^36^. BMH-21 is a DNA intercalator that inhibits rRNA synthesis, promoting the disassembly of the polymerase from the rDNA and the subsequent degradation of Rpa190 ^36, 37^. Interestingly, *rpa190-KR* mutant cells were more resistant to BMH-21 (Figure 3F). This phenotype was dominant, as expression of an *rpa190-KR* allele from a centromeric vector in wild type cells conferred resistance to BMH-21 (Figure 3G). In addition, *ubp10Δ* mutant cells were more sensitive to BMH- 21, as expected from their increased Rpa190 ubiquitination levels. Moreover, ectopic expression of an *rpa190-KR* allele rescued the BMH-21 sensitivity of *ubp10Δ* mutant cells, evidencing that K408 and K410 play a pivotal role in the response to BMH-21. Overall, our findings indicate that Rpa190 ubiquitination at K408 and K410 regulates RNAP I under conditions that prevent transcriptional elongation.

### An active Smc5/6 complex is required to sustain normal Rpa190 ubiquitination levels

To validate the decrease in Rpa190 ubiquitination in *nse1-RING* mutants, we pulled down 7His-tagged ubiquitin in cells expressing a 6HA-tagged copy of Rpa190. Ubiquitination levels of Rpa190 were lower in *nse1-C274A*, relative to wild type cells (Figure 4A), indicating that this modification is partly dependent on the Nse1 RING domain. The reduction was even more evident after treatment with MMS, conditions that increased Rpa190 ubiquitination in wild type cells. In addition, the Rpa190 ubiquitination impairment was similar in *nse1-C274A* and *rpa190-KR* cells (Supplementary Figure 5). To analyze the participation of the Smc5/6 complex in Rpa190 ubiquitination, we tested various hypomorphic Smc5/6 mutants. As shown in Figure 3B, *nse5-ts3* and *nse1-16*, displayed reduced Rpa190 ubiquitination levels, indicating that a fully functional Smc5/6 complex is required to sustain normal Rpa190 ubiquitination.

**Figure 4.**
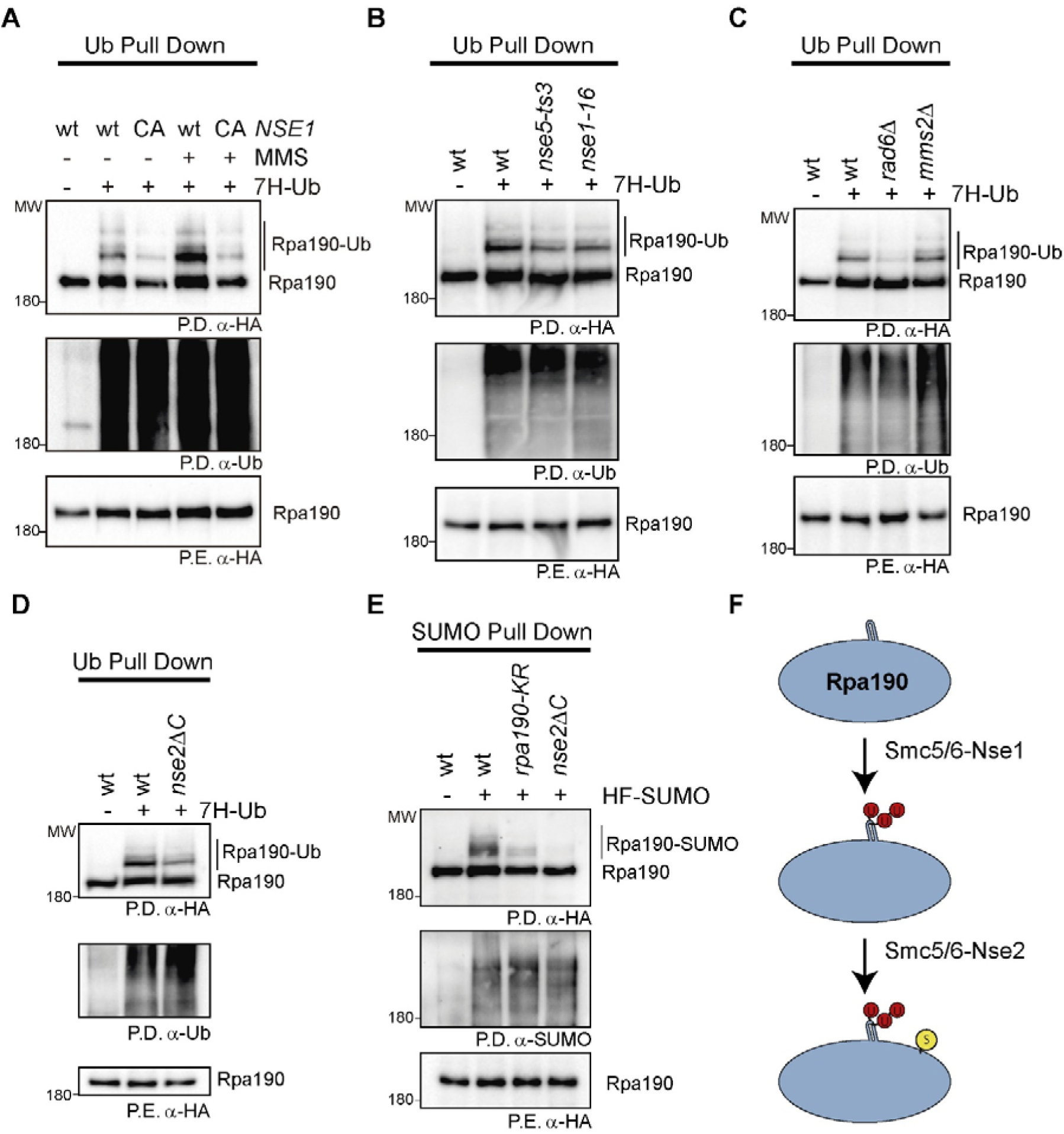
An active Smc5/6 complex is required to sustain normal Rpa190 ubiquitination levels. **A.** Pull down analysis of Rpa190 ubiquitination in wild type (wt) and *nse1-CA* cells; ubiquitin was pulled down (P.D.) under denaturing conditions from yeast protein extracts (P.E.) to purify ubiquitinated species. **B.** Denaturing ubiquitin pull down analysis in wild type, *nse5-ts3* and *nse1-16* cells grown at the permissive temperature (25°C). **C.** Denaturing ubiquitin pull down analysis of Rpa190 ubiquitination in wild type and E2 (*rad6Δ* or *mms2Δ*) mutant strains. **D.** Denaturing ubiquitin pull down analysis of Rpa190 ubiquitination in wild type and *nse2ΔC* mutant strains. **E.** Denaturing SUMO pull down analysis of Rpa190 SUMOylation in *rpa190-KR* and *nse2ΔC* mutant strains. **F.** Proposed model for sequential modification of Rpa190 by Nse1-dependent ubiquitination at K408 and K410 (U, in red) followed by Nse2-dependent SUMOylation (S, in yellow). In A-E, 6xHA-tagged Rpa190 was detected with anti-HA antibodies; ubiquitin or SUMO-conjugated forms of Rpa190 are indicated with a vertical bar.

Monoubiquitination of Pol30 depends on Rad6, while the Ubc13-Mms2 E2 enzymes stimulate its polyubiquitination. Since Pol30 ubiquitination is reduced in *nse1-C274A* cells, we tested if these E2s also participate in modification of Rpa190. As shown in Figure 4C, Rpa190 ubiquitination levels were lower in *rad6Δ* cells, while they were not affected by the *mms2Δ* mutation, suggesting that Rad6 promotes ubiquitination of Rpa190.

Interestingly, SUMOylation of RNAP I subunits is controlled by the Nse2 subunit in the Smc5/6 complex ^39^. Thus, Rpa190 SUMOylation and ubiquitination might be interdependent. To analyze this possibility, we tested Rpa190 ubiquitination in *nse2ΔC* mutant cells, which are severely impaired in Rpa190 SUMOylation (Figure 4E). Interestingly, Rpa190 ubiquitination was diminished in *nse2ΔC* mutants, relative to wild type cells (Figure 4D). This suggests that perturbing RNAP I SUMOylation might affect its subsequent ubiquitination; alternatively, the reduced Rpa190 ubiquitination might derive from deficient Smc5/6 function in *nse2ΔC* cells, as occurs in *nse5-ts3* and *nse1-16* mutants (Figure 4B). In contrast, Rpa190 SUMOylation was severely impaired in *rpa190-KR* cells (Figure 3E), suggesting that Rpa190 ubiquitination is required for its subsequent SUMOylation (Figure 4F). Of note, despite the reduced SUMOylation of the Rpa190-KR protein, the pattern of SUMO bands was similar to that of wild type Rpa190 (Figure 4E), suggesting that SUMO may not target K408 and K410. Overall, our results indicate that Smc5/6 function is required to sustain proper Rpa190 ubiquitination levels which, in turn, promotes its subsequent modification with SUMO.

### Rpa190 ubiquitination improves rDNA segregation in *smc5/6* mutants

We have previously demonstrated that RNAP I function is detrimental for rDNA segregation in *smc5/6* mutants ^8^. Therefore, it is possible that Smc5/6 might promote local clearance of stalled RNAP I complexes through ubiquitination of the Rpa190 subunit, thus ensuring completion of rDNA replication and sister chromatid disjunction. To test this idea, we analysed rDNA segregation, using a battery of tet operators (tetO:487) inserted in the telomeric flank of the rDNA array ^40^. Cultures of *smc5/6* mutants cells typically fail to segregate tetO:487 in 50% of the population ^29^. To inactivate Smc5/6 function, we used an auxin inducible degron allele of *SMC5* (*smc5-aid*). After synchronization of cells in G1 with alpha factor, cultures were treated with auxin to promote Smc5-aid degradation. Next, cells were released from the arrest and observed under the microscope. Cells started budding at 40 minutes, entered anaphase between 80 and 100 minutes, and most of them completed nuclear fission 140 minutes after release from G1 (Figure 5A). Cells with two divided nuclei at time points between 120 and 160 minutes were scored for segregation of tetO:487 operators (Figure 5B). As shown in Figure 5C, the *rpa190-KR* allele significantly aggravates the rDNA segregation defect of *smc5-aid* cells. In contrast, deletion of *UBP10* had the opposite effect and improved rDNA segregation in *smc5- aid* cells. Therefore, our results indicate that Rpa190 ubiquitination levels affect rDNA segregation in *smc5-aid* mutant cells and suggest that rDNA missegregation in *smc5/6* mutant cells partly stem from insufficient Rpa190 ubiquitination. Overall, these observations suggest that Smc5/6-dependent ubiquitination of Rpa190 might be a mechanism to dislodge stalled RNAP I complexes from chromatin, thus preventing non-disjunction events in the rDNA array.

**Figure 5.**
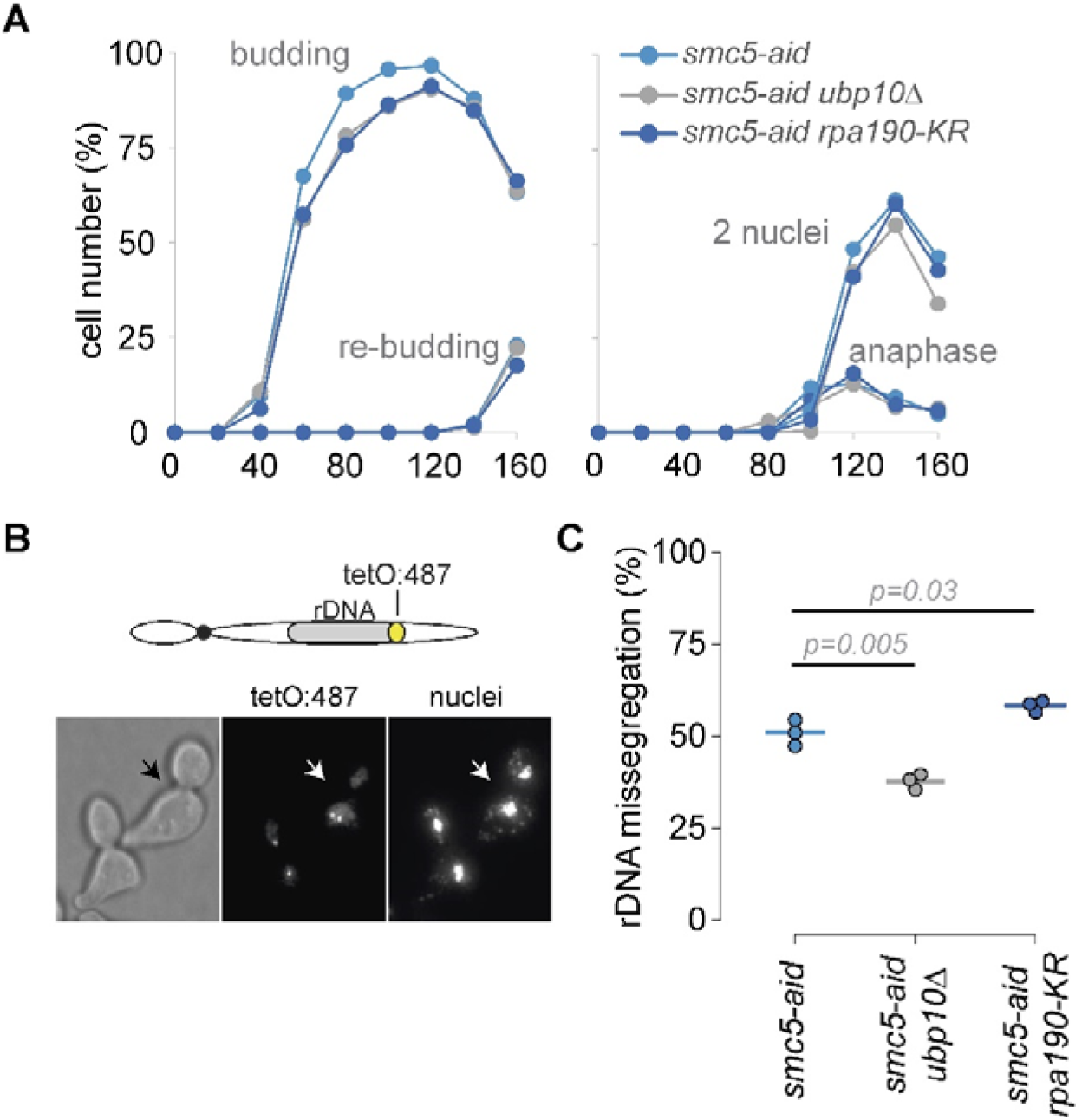
Rpa190 ubiquitination improves rDNA segregation in *smc5/6* mutants. **A.** Cells of the indicated genotype were synchronized in G1 with alpha factor, treated with auxin for 30 min to degrade Smc5-aid and released into a synchronous cell cycle in the presence of auxin. Samples were taken at the indicated time points for microscopic analysis and scored for budding (first S phase), re-budding (second S phase), elongated nucleus between mother and daughter cell (anaphase) and 2 nuclei (two nuclear masses between mother cell and bud). **B.** Around 50% of *smc5-aid* fail to segregate the distal part of the rDNA locus; upper panel, rDNA segregation was assayed using a battery of fluorescent *tetO* repeats inserted at the indicated position in chromosome XII; bottom panel, representative micrographs of *smc5-aid* cells at time 120 min after G1 release, nuclei stained with Hoetsch; arrow points to cell with missegregated *tetO* tags. **C.** Quantification of rDNA missegregation in cells of the indicated genotypes. Cells with two nuclei at times 105-160 min after G1 release were scored for rDNA segregation, as shown in B. Circles indicate individual measurements from 3 independent experiments; horizontal lines show medians; *p*=*p*-value, ANOVA test.

## DISCUSSION

### A proteomic approach to identify Nse1-dependent ubiquitin targets

The notion that inactivation of the RING domain in Nse1 impairs cell growth and sensitizes cells to DNA damage highlights its important functions in cell proliferation and genome integrity ^17, 21, 22^. Nse1 has ubiquitin E3 ligase activity *in vitro*. This activity is enhanced by the Nse3 subunit ^14^. However, the only *in vivo* substrates identified for Nse1 are (i) the Nse4 subunit of the Smc5/6 complex in yeast ^19^ and (ii) Mms19, a cytosolic iron-sulfur assembly (CIA) component, in humans ^20^. Our label-free proteomic approach has allowed us to identify a large set of Nse1-dependent ubiquitin targets, and to reveal novel and important connections of Nse1 with DNA replication and DNA damage tolerance, ribosomal biogenesis and cell metabolism. Thus, our study should constitute a valuable source of information about the ubiquitin-dependent processes and pathways affected by the RING domain in Nse1.

Network analysis of Nse1-dependent ubiquitin targets exposes several functional clusters (Figure 1). Interestingly, a large set of proteins, including Pol I subunits, RNA processing elements and ribosomal proteins, show deregulated ubiquitination in *nse1* mutants cells, what points to ribosome biogenesis as one of the main processes controlled by Nse1. This is in accordance with the previously reported nuclear accumulation of ribosomal proteins and the ribosome biogenesis defects in *smc5/6* and *nse2ΔRING* mutants ^41^, which could result from deficient controls during ribosome biogenesis. Given the relevance of ribosome biogenesis for optimal cell growth and proliferation, it is possible that the impaired fitness of *nse1* mutant cells ^21, 22^ may be partly due to defects in ribosome synthesis and altered protein translation.

Another notable ubiquitin target identified in our proteomic screen is K164 in Pol30 (PCNA). PCNA is known to be monoubiquitinated at K164 by Rad6-Rad18 and further polyubiquitinated with K63-linked ubiquitin chains by Mms2-Ubc13-Rad5 ^42, 43^. In vitro, PCNA ubiquitination is very inefficient and requires prior RFC-dependent loading of PCNA onto partially heteroduplex DNA ^44, 45^. The requirements may be even more intricate in vivo, as PCNA ubiquitination occurs in a context of damaged and partially replicated chromatin fibers. We envisage that Smc5/6-Nse1 may coordinate sister chromatid organization and PCNA modification behind forks to enable efficient bypass of DNA lesions. In accordance, the Smc5/6 complex is recruited to damaged forks, where it aids in replisome stability and fork remodeling ^46, 47^. Notably, a direct link between the Smc5/6 complex and the PCNA ubiquitination machinery has been previously reported in vertebrates: Rad18 interacts with the Nse5-Nse6 homologs to recruit the Smc5/6 complex to interstrand crosslinks, a burdensome lesion that stalls replication forks ^48^. However, we cannot discard the possibility that Nse1 plays an ancillary role in Pol30 ubiquitination, perhaps by reconfiguring the topology of damaged replication forks, in an unknown way, to prime full PCNA ubiquitination.

Other groups of ubiquitin targets identified in our study are related to glycolytic or sterol synthesis processes that, apparently, are not directly related to Smc5/6 or Nse1 functions on chromatin. Interestingly, some mutants in the ergosterol biosynthesis are sensitive to DNA damage, suggesting that one or more metabolite/s in this pathway could play a role in DNA damage repair ^49^. On the other hand, it is worth noting that previous studies have also established a connection between the Smc5/6 complex, DNA damage and carbon metabolism ^50^. Moreover, hypomorphic *NSMCE2* mutations identified in two patients have been related to a human syndrome linked to dwarfism, insulin resistance and severe metabolic abnormalities^51^. Based on our proteomic data, we propose that the Smc5/6-dependent regulation of metabolic processes may be channeled, at least partly, through protein ubiquitination.

### A role for Nse1-dependent ubiquitination in Pol I transcriptional elongation

Another ubiquitin target identified in our screen is Rpa190, the large subunit of RNAP I. Based on our proteomics and pull-down experiments, we propose that ubiquitination of lysines 408 and 410 depends on Nse1. Of note, the Smc5/6 complex and RNAP I show some interesting connections. First, both of them bind to the rDNA array ^29, 52^; besides, RNAP I is SUMOylated in an Nse2-dependent manner ^39^; in addition, inactivation of RNAP I partially relieves the rDNA missegregation phenotype of *smc5/6* mutants ^8^.

Ubiquitination levels of two Nse1-dependent ubiquitin targets identified in this study, Rpa190 and Pol30, seem to be regulated by similar players, as both depend on Rad6 but also on the Ubp10 ubiquitin protease ^34, 53^. However, the outcome of the modification might be different in each target: Rpa190 ubiquitination affects its stability, while Pol30 ubiquitination does not. Our findings also suggest that stalling of elongating RNAP I complexes by UV light damage induces Rpa190 ubiquitination. In turn, Rpa190 ubiquitination can promote its degradation by the proteasome, a situation that is normally counteracted by Ubp10. The analysis of double *rpa190-KR ubp10Δ* mutants indicates that K408 and K410 are the relevant targets for ubiquitin-dependent Rpa190 degradation. We speculate that cells may ubiquitinate stalled RNAP I complexes to degrade them and prevent further interferences with other transactions at the rDNA locus. As ongoing RNAP I complexes run into DNA replication and repair processes, Nse1 dependent-ubiquitination may promote RNAP I clearance from individual rDNA repeats and allow resolution of sister rDNAs. In accordance, we have shown that preventing Rpa190 deubiquitination worsens rDNA missegregation in Smc5/6 mutant cells, while the *rpa190-KR* mutant improves it. Thus, Rpa190 ubiquitination facilitates disjunction of the rDNA array, helping to maintain its integrity.

The localization of the ubiquitination sites in Rpa190, at the tip of the clamp domain (Figure 3A), might shed clues about the role of this modification. During initiation of transcription, the cleft aperture in RNAP I changes from an open conformation when bound to the Rrn3 transcription factor, to a fully closed configuration in the elongating complex ^54, 55^. In addition, the Rpa49 linker crosses on top of the cleft, passes over the downstream rDNA, and binds just behind the clamp domain, acting as a safety belt and securing the rDNA inside the transcribing RNAP I complex. Ubiquitination sites in Rpa190 lay close to the path followed by the Rpa49 linker domain, whose roles seem to be the stable clamping of the rDNA during transcriptional elongation. Thus, we hypothesize that ubiquitination at the tip of the clamp might interfere with Rpa49, perhaps reducing the processivity of RNAP I in preparation for its unloading from the rDNA. Sequence and structure alignment indicates that lysine 350 in the RPA194 subunit of the human RNAP I complex occupies the same position as lysine 408 in yeast (Supplementary Figure 3); moreover, quantitative diGly proteomics has shown that K350 in RPA194 is also modified in vivo ^23^, suggesting that this mechanism might be conserved in humans.

Although UV damage-induced Rpa190 ubiquitination is not accompanied by a global reduction in Rpa190 protein levels, it is probable that Rpa190 ubiquitination occurs locally, to remodel RNAP I complexes stalled by UV damage and facilitate DNA repair, in an analogous way to RNAP II ubiquitination during transcription coupled nucleotide excision repair (TC-NER) ^56^. In accordance, RNAP I density along rDNA units is altered after UV-light damage, most probably as a result of its dissociation from the central and 3’-terminal side of the transcriptional unit ^57^. One intriguing aspect of our study is that *rpa190-KR* mutants are generally healthy and do not exhibit other phenotypes in common with Smc5/6 mutants, such as sensitivity to DNA damage, including lesions inflicted by UV light. However, while RNAP II ubiquitin mutants are not UV sensitive, they aggravate the sensitivity of TC-NER mutants ^58^. Thus, onsite RNAP I ubiquitination might be used as a backup resort when conventional DNA repair pathways fail. Interestingly, we have identified other ubiquitin sites in RNAP I significantly under-ubiquitinated in *nse1-C274A* mutant cells. For example, Rpc19, a subunit shared by RNAP I and RNAP III, showed lower ubiquitination on two different sites (with 3.5 and 2.6 fold reduction) in *nse1* mutant, relative to wild type. In comparison, Rpa190 ubiquitination at both K408 and K410 was reduced 2.8 fold in *nse1* mutant extracts. In addition, K120 in Rpa34 and K88 in Rpa43 were also less ubiquitinated in *nse1* mutants. Moreover, our proteomic screen allowed us to identify five additional sites under-ubiquitinated in *nse1* mutant cells (two sites in Rpa190 and one in Rpa135, Rpb5 and Rpb10), although differences were not statistically significant, relative to the wild type. Therefore, if ubiquitin redundantly targets multiple lysines in RNAP I, the modification of other sites in the complex may be masking a more severe phenotype in *rpa190- K408,410R* cells. It will be interesting to extend RNAP I KR mutations to other Nse1- dependent sites, and test how it relates to Smc5/6-dependent functions.

Cancer cells frequently increase rRNA expression, thus making RNAP I as an attractive candidate for cancer treatment ^59^. Remarkably, *rpa190-KR* mutants are resistant to BMH-21, a specific transcriptional elongation inhibitor of RNAP I. The resistance of *rpa190-KR* cells to BMH-21 opens the question about what is the nature of the problem generated by this inhibitor. BMH-21 is believed to bind G-C rich areas at the rDNA and to block transcriptional elongation by RNAP I ^36, 37^. The resistance of non-ubiquitinable *rpa190-KR* mutants suggests that the sensitivity to BMH-21 is primarily due to Rpa190 ubiquitination, and not to the inability to proceed with transcriptional elongation. As yeast and human cells are equipped with similar tools to regulate Rpa190 ubiquitination, analysis of the physiological situations and mechanisms that promote RNAP I ubiquitination in cancer cells will be of great interest.

## METHODS

### Yeast strains

All strains used in this study are derivatives of EMY74.7 (*MATa his3-Δ1 leu2-3,112 trp1Δ ura3-52*), W303 (*MATa ade2-1 can1-100 his3-11,15 leu2-3,112 trp1-1 ura3-1 RAD5+*) or BY4741 (*MATa his3Δ1 leu2Δ0 met15Δ0 ura3Δ0*). A list of yeast strains used in this study, together with their relevant genotype, the plasmids they carry and the figure where they are used, is provided in Supplementary Tables S1 and S2. Yeast cells were grown in YP (Yeast extract and Peptone) or synthetic complete (SC) drop-out medium for selection of expression plasmids, plus 2% glucose. For auxin-induced degrons (*smc5-aid*), IAA (SIGMA) was added to 1 mM from a 0,5 M stock in water.

### Ubiquitinome profiling

#### Protein digestion

To identify Nse1-dependent ubiquitin targets, we isolated the nuclear fraction from 2 l of wild type and *nse1-C274A* mutant yeast cells treated with MMS 0.02% for 1,5 hours, until reaching an OD_600_∼1.5. Cells were harvested for 5 min at 5.000 rpm, resuspended and incubated for 5 min in 400 ml of azide solution (50 mM Tris-Cl-H pH 7.5, azide 0.1% (15 mM)). Next, cells were centrifuged for 5 min at 5.000 rpm, resuspended in 400 ml of reduction solution (500 µl of 1M Tris-HCl pH 7.5, 250 µl β-ME, 49,25 ml H_2_O) and incubated 30 min at room temperature. Cells were pelleted and resuspended in 100 ml of SP buffer (1 M sorbitol, 20 mM phosphate buffer pH 7.4) and 2 mg/gr of cells of zymoliase 100T was added to each culture. Then, samples were incubated at 25°C for 20-25 min at a 100 rpm shaking agitation. After checking spheroplasting, cells were sedimented at 3.200 rpm for 3 min and washed with SP buffer. After removing the supernatant, spheroplasts were resuspended in Ficoll buffer (18% FICOLL, 20 mM phosphate buffer pH 6.8 (PI-Roche, 1 mM EDTA, 5 mM chloroacetamide, 50 µM PR-619)) and kept on ice for 5 min. Samples were centrifuged at 30.000 g for 20 min at 4°C. Nuclear pellets were lysed in 10 volumes of lysis buffer (5% Sodium deoxycholate, 100 mM Tris, pH 8) supplemented with protease and phosphatase inhibitors (Halt, Pierce). The lysate was incubated at 95°C 10 minutes and sonicated (20% intensity, 2 min, 1 sec on, 1 sec off). After cooling, 1:1000 (v/v) Benzonase (Merck) was added and further incubated for 1 hour at 37 C. Proteins were reduced and alkylated for 1 h with 15 mM TCEP (Sigma) and 30 mM CAA (Sigma) at RT in the dark. Lysates were diluted 1:5 with 50 mM Tris HCl, pH 8, and proteins were digested with Lys-C (Wako) (enzyme: substrate ratio 1:50) for 6 h. Subsequently, samples were digested overnight at 37 °C with recombinant trypsin (Trypzean, Sigma) (enzyme: substrate ratio 1:50). Enzymes activity were stopped with 2% TFA (v/v). Deoxycholic acid was removed by centrifugal precipitation (20 minutes at 20,000 x g). Peptides present in the supernatant were desalted with 500 mg C18 SepPak SPE cartridges (Waters) and lyophilized.

#### Immunoprecipitation of di-Gly–containing peptides

Enrichment of K-ε-GG peptides was performed with the PTMScan ubiquitin remnant motif (K-ε-GG) kit (Cell Signaling Technology, cat. no. 5562) containing anti–K-ε-GG antibody cross-linked to protein A beads. Ten milligrams of lyophilized peptides per condition were gently resuspended in 1200 μl of ice cold immunoaffinity purification (IAP) buffer (50 mM MOPS, pH 7.4, 10 mM Na2HPO4, and 50 mM NaCl) and centrifuged at maximal speed for 5 min at 4°C to remove insoluble materials. The supernatants were mixed with 20 μl of anti–K-ε-GG of pre-conditioned antibody beads and incubated for 2 h at 4°C with rotation. Samples were centrifuged and the supernatant containing the unbound fraction was used to determine the total protein content. Beads were washed twice with 1 ml of ice-cold IAP buffer followed by three washes with ice-cold PBS and two washes with ice-cold MS-grade water. K-ε-GG peptides were eluted from the antibody with 2 × 50 μl of 1% TFA. Eluted peptides were desalted with C18 StageTips conditioned by washing with 80 μl of methanol followed by an equilibration with 80 μl of 1% TFA. Samples were loaded on StageTips, washed twice with 80 μl of 0.1% formic acid FA and eluted with 50 μl 60% MeCN/0.1% (FA). Eluted peptides were dried completely with vacuum centrifugation and reconstituted in 0.1% FA.

#### Mass spectrometry

LC-MS/MS was carried out by coupling an Ultimate 3000 RSLCnano System (Dionex) with a Q-Exactive HF-X mass spectrometer (ThermoScientific). Samples were loaded into a trap column (Acclaim PepMap^TM^ 100, 100 µm x 2 cm, ThermoScientific) over 3 min at a flow rate of 10 µl/min in 0.1% FA. Then peptides were transferred to an analytical column (PepMap^TM^ RSLC C18, 2 µm, 75 µm x 50 cm, ThermoScientific). The fractions containing di-Gly-containing peptides were separated using a 60 min effective linear gradient (buffer A: 0.1% FA; buffer B: 100% ACN, 0.1% FA) at a flow rate of 250 nL/min. The gradient used was: 0-3 min 2% B, 3-5 min 6% B, 5-36 min 17.5% B, 36-60 min 25% B, 60-63 min 33% B, 63-65 min 45% B, 65-70 min 98% B, 70-80 min 2% B. The peptides were electrosprayed (1.9 kV) into the mass spectrometer through a heated capillary at 300 °C and an ion funnel RF level of 40%. The mass spectrometer was operated in a data-dependent mode, with an automatic switch between the MS and MS/MS scans using a top 12 method (minimum AGC target 8E3) and a dynamic exclusion time of 25 sec. MS (350-1500 m/z) and MS/MS spectra were acquired with a resolution of 60,000 and 30,000 FWHM (200 m/z), respectively. Peptides were isolated using a 2 Th window and fragmented using higher-energy collisional dissociation (HCD) at 27% normalized collision energy. The ion target values were 3E6 for MS (25 ms maximum injection time) and 1E5 for MS/MS (54 ms maximum injection time). Two independent biological replicates and two technical replicates were analyzed.

The peptides contained in the unbound fraction were separated using a 90 min effective linear gradient (buffer A: 0.1% FA; buffer B: 100% ACN, 0.1% FA) at a flow rate of 250 nL/min. The gradient used was: 0-5 min 4% B, 5-7.5 min 6% B, 7.5-80 min 25% B, 80-94 min 42.5% B, 94-99 min 98% B, 99-105 min 2% B. The peptides were electrosprayed (1.9 kV) into the mass spectrometer through a heated capillary at 300 °C and an ion funnel RF level of 40%. The mass spectrometer was operated in a data-dependent mode, with an automatic switch between the MS and MS/MS scans using a top 12 method (minimum AGC target 8E3) and a dynamic exclusion time of 40 sec. MS (350-1500 m/z) and MS/MS spectra were acquired with a resolution of 60,000 and 15,000 FWHM (200 m/z), respectively. Peptides were isolated using a 2 Th window and fragmented using higher-energy collisional dissociation (HCD) at 27% normalized collision energy. The ion target values were 3E6 for MS (25 ms maximum injection time) and 1E5 for MS/MS (22 ms maximum injection time). Samples were analyzed twice.

#### Data analysis

Raw files from both K-ε-GG–enriched fraction and full proteome (flow-through, unbound fraction) were processed with MaxQuant (version 1.6.0.16) using the standard settings against a *Saccharomyces cerevisiae* protein database (UniProtKB/Swiss-Prot, 6,718 sequences) supplemented with contaminants. Label-free quantification was done with match between runs (match window of 0.7 min and alignment window of 20 min). Carbamidomethylation of cysteines was set as a fixed modification whereas addition of glycine-glycine to lysine (only for K-ε-GG–enriched fraction), methionine oxidation and N-term acetylation were variable protein modifications. The minimal peptide length was set to 7 amino acids and a maximum of two tryptic missed-cleavages were allowed. The results were filtered at 0.01 FDR (peptide and protein level) and subsequently the “proteinGroup.txt” file and the “GlyGly (K) Sites.txt” were loaded in Perseus (v1.6.0.7) for further statistical analysis. The intensity of the K-ε-GG–enriched fraction was normalized by subtracting the median. A minimum of 75% valid values per group was required for quantification. Missing values were imputed with Perseus from the observed normal distribution of intensities. Then, a two-sample Student’s T-Test with a permutation-based FDR was performed. Proteins with a q-value<0.05 and log_2_ ratio >0.5 or < -0.5 and K-ε-GG– peptides with a q-value<0.1 and log_2_ ratio >1 or < -1 were considered as regulated. The mass spectrometry proteomics data have been deposited to the ProteomeXchange Consortium via the PRIDE partner repository with the dataset identifier PXD028669.

### Ubiquitin and SUMO Pull downs

Ubiquitin and SUMO pull-downs were done under denaturing conditions using protein extracts from yeast cells carrying the *UBI* gene fused to a 7xHis tag, expressed from a plasmid under the control of the *CUP1* promoter; or the *SMT3* gene fused to a 6xHis-Flag epitope integrated at its endogenous location. For *UBI* expression from the *CUP1* promoter, cultures were grown in SC-URA medium containing CuSO_4_ at 20 µM.

25 OD_600_ of exponentially growing cells (100 OD_600_ for SUMOyated Rpa190 PD) were spun at 3.000 rpm 2 minutes in 50 ml tubes. Cells were then washed in cold water and transferred to a 1.5 ml tube. 250 µl of Buffer A (6 M Guanidine Chloride, 100 mM KH_2_PO_4_, 10 mM Tris-HCl, 0.05% Tween-20, 15 mM imidazole, 1X protease inhibitors; pH 8) and 500 µl of glass beads were added to the pellet. Cells were broken using a mini-beadbeater cell disrupter (BioSpec Products) at power 6 for 1,5 minutes. Next, we pierced the bottom of the tube with a 21 ga. needle and set it up over another 1.5 ml tube. After that, extracts were spun at 2.400 rpm to recover the cell extract and discard the beads. Following, we added 700 µl of Buffer A to the tube, mixed it and spun 10 minutes at 14.000 rpm at 4°C. The supernatant was then transferred to a screw cap tube, and the volume brought up to 1 ml. Finally, we added 70 µl of Ni-NTA (Nickel Nitrilo-triacetic Acid) agarose beads (Qiagen), and incubated the mixture overnight at the orbital rotator at RT. Next, the tubes were spun at 3.400 rpm. The supernatant was transferred to a new tube, and the beads washed for 8 minutes with 900 µl of Buffer B (8 M urea, 100 mM KH_2_PO_4_, 10 mM Tris-HCl, 0.05% Tween-20, 2 mM imidazole; pH 8). Washes were repeated a total of three times with Buffer B and three times more with Buffer C (8 M urea, 100 mM KH_2_PO_4_, 10 mM Tris-HCl, 0.05% Tween-20; pH 6.3). Finally, proteins bound to the beads were eluted with 25 µl of 2xSSR (4% SDS, 250 mM Tris-HCl pH 6.8, 10% sucrose, 0.02% bromophenol blue) by boiling 3 minutes at 95°C. 30 µl of supernatants were precipitated with the same volume of TCA 12% (trichloroacetic acid), washed 3 times with chilled acetone, resuspended by vortexing in 30 µl of Buffer C and incubated at 37°C for 20 min. Then, protein extracts (PE) were boiled with 30 µl of 2xSSR. Finally, PE and PDs were centrifuged and loaded onto a SDS-PAGE for Western Blot analysis.

### Western blot analysis

This protocol was used for analysing Pol30-protein levels and modifications by Western Blot. Briefly, 10 OD_600_ from an exponentially growing culture were collected by centrifugation 4 minutes at 4.000 rpm and 4°C, after adding 250 µl of TCA 20% directly to the culture. The pellet was resuspended with 500 µl of TCA 20%, transferred to a 1.5 ml tube and spun 6-7 seconds at high speed. Then, the supernatant was removed and the pellet was resuspended again with 100 µl of TCA 20%. After that, we added 500 µl of glass beads and broke cells in a mini-beadbeater cell disrupter (BioSpec Products) for 45 seconds. Next, the sample was transferred to a new tube by making a hole with a needle at the bottom and spinning 1 minute at 1.000 rpm. After that, we washed beads twice with 100 µl of TCA 5% and centrifuged 10 minutes at 3.000 rpm and RT. The supernatant was discarded and the pellet was resuspended with 100 µl of 1xSR (2% SDS in 0.125 M Tris-HCl pH 6.8) and neutralized with 50 µl of Tris-base 1 M. Samples were then boiled 3 minutes at 95°C, centrifuged 5 minutes at 13.000 rpm and protein concentration was determined with the BIO-RAD Micro DC Protein Assay. Finally, 4xSS and β-Mercaptoethanol were added to a volume corresponding to 30 µg of protein, loaded onto an SDS-gel and analyzed by Western Blot.

### Generation of *rpa190-KR* mutants

To obtain the *rpa190-K408,410R* allele, we mutagenized a plasmid expressing the wild type gene (*pRS314-(CEN-TRP1*)-*RPA190*) by PCR using a pair of primers containing the two point mutations and a PstI restriction site. Colonies were checked by PstI digestion and sequencing. The mutant and the wild type plasmids were then transformed into a diploid yeast strain with a deletion in one of the endogenous copies of *RPA190*. Haploid yeast colonies carrying the *rpa190-K408,410R* plasmid and a deleted chromosomal *RPA190* copy were isolated by sporulation and tetrad dissection. Next, we tagged the plasmid express copy with a 6HA tag and recovered the vectors from their yeast to obtain the *pRS314-(CEN-TRP1)-RPA190- 6HA:natNT2* and *pRS314-(CEN-TRP1)-rpa190-K408,410R*-6HA:natNT2 plasmids. To integrate the double *KR* mutation into yeast, we used the mutant vectors as PCR templates. The PCR product was transformed into yeast and colonies carrying the desired mutation were selected by PstI restriction and sequencing.

### Gene tagging

Gene tagging was performed as previously described ^60^, using a PCR-based method for recombinational integration. A set of cassette plasmids carrying a selectable marker and an epitope tag were amplified by PCR using the adequate primers to direct the insertion of the cassette to the desired locus by homologous recombination.

### Yeast growth test analysis

10-fold dilution series of yeast cultures were spotted on solid medium. Plates were incubated at the adequate temperature during 2-3 days and photographed with the Chemiluminiscent Imager (Bio-Rad). For methyl methanesulfonate (MMS) growing test analysis, MMS was added from 0.001 to 0.02% final concentrations in the YPD agar media before solidification. BMH-21 (Sigma) was added to the YPD agar at a 15 µM concentration. Plates were incubated at 30°C until individual colonies became visible in control plates.

### Cell Cycle experiments

Exponentially growing cells were arrested in G1 by addition of 10^-6^ M alpha factor (Genscript), in G2/M by addition of 15 μg/ml nocodazole (SIGMA) or 0.2 M hydroxyurea (SIGMA) at 25°C for 2 hours. In Figure 5, cultures were arrested in G1 by addition of 10^-8^ M alpha factor, and released by washing cells 3 times with pre-warmed medium and re-suspended in media containing 0,1 mg/ml pronase E (SIGMA). For microscopic visualization, DNA was stained using Hoechst at 0,5 µg/ml final concentration in the presence of mounting solution and 0,4% Triton X- 100 to permeabilize cells. For fluorescence microscopy of nuclei or tetO:487 dots, series of z-focal plane images were collected with a DP30 monochrome camera mounted on an upright BX51 Olympus fluorescence microscope.

## AKNOWLEDGEMENTS

The work in JT-R lab was supported by grants BFU2015-71308-P and PGC2018- 097796-B-I00 from the Ministerio de Ciencia, Innovación y Universidades, and grant 2017-SGR-569 from AGAUR-Generalitat de Catalunya; the IRBLLEIDA institute is part of the CERCA Programme/Generalitat de Catalunya. The CNIO Proteomics Unit belongs to ProteoRed, PRB3- ISCIII, supported by grant PT17/0019/0005. Eva Ibars was supported by a PhD fellowship from UdL and ‘Ajuts al talent en investigació Biomèdica’ contract from IRBLLEIDA and Diputació de Lleida. We thank Farida Dakterzada for construction of yeast strains, Paul Kaufman for the polyclonal anti- Pol30 antibody, Herbert Tschochner for the *pRS314-RPA190* plasmid, Rodrigo Bermejo for the *YEplac195-CUP1p-7xHis-UBI* plasmid and Carlos Fernéndez- Tornero and all members of the Cell Cycle lab for helpful discussions.

## Author contributions

E.I., J.C.F., G.B., C.C., M.T., R.S-S., N.P.L., N.C. and J.T-R. performed the experiments. P.X-E. and J.M. performed ubiquitin profiling. E.I., J.C.F., G.B. and J.T-R. analysed and interpreted the data. J.T-R. designed the project and wrote the manuscript. All authors read and approved the final version of the manuscript.

## Competing interests

The authors declare no competing interests.

---

**Supplementary Figure 1.**
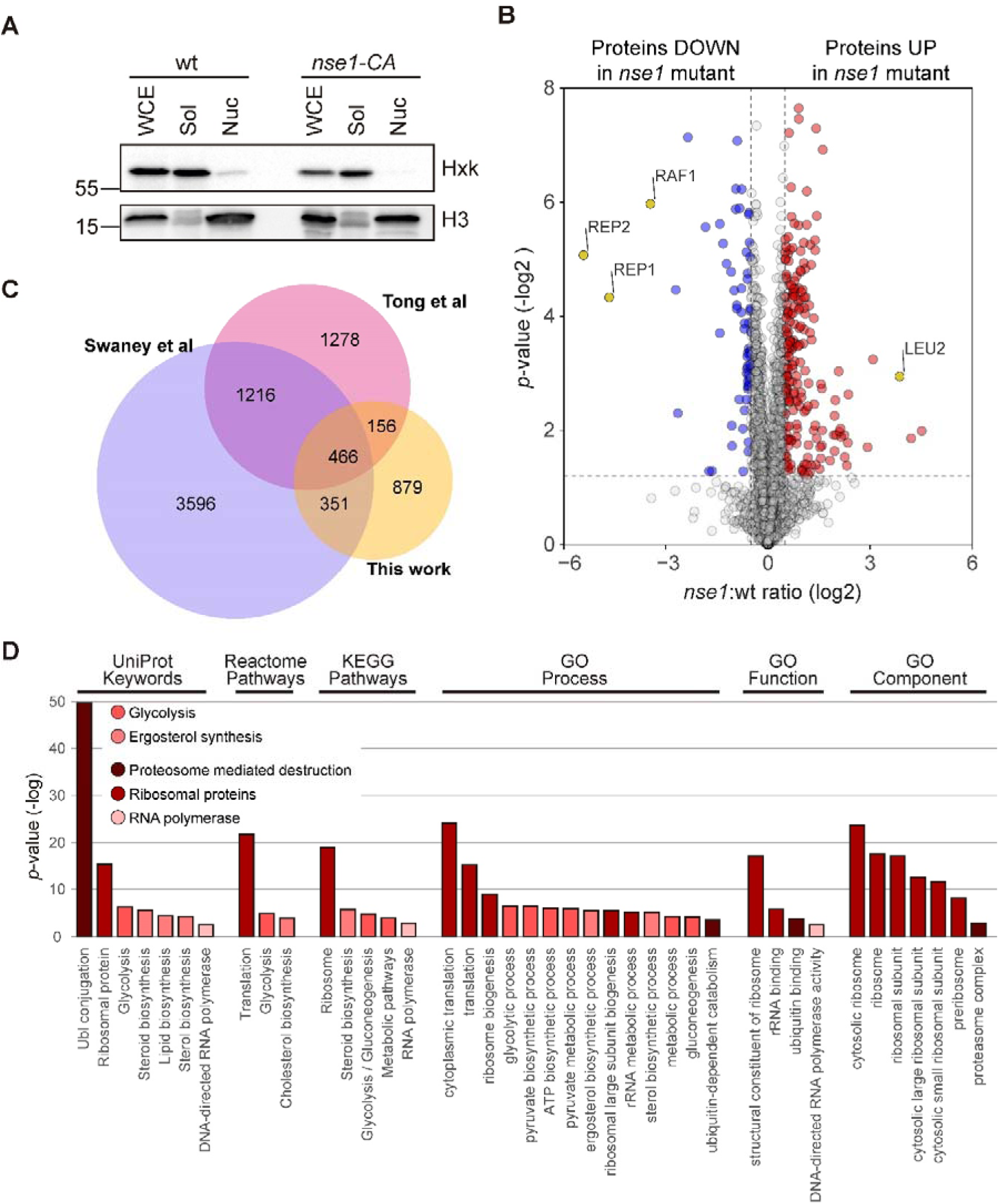
A proteomic screen to identify Nse1-sensitive ubiquitin sites. **A.** Fractionation of whole cell extracts (WCE) from wild type (wt) and *nse1-C274A* (*nse1-CA*) mutant yeast cells treated with MMS 0.02% into soluble cytosolic (Sol) and nuclear (Nuc) fractions. Protein samples from WCE, Sol and Nuc were analysed by western blot and tested for the presence of hexokinase (Hxk), a soluble protein, and histone H3 (H3), a nuclear protein. **B**. Volcano plot showing the differentially expressed proteins (log_2_ ratios) between wild type and *nse1-C274A* mutant cells on the x-axis and their statistical significance (−log_10_(*p*-value)) on the y-axis; significantly affected proteins are shown above the dotted line; proteins down- regulated in *nse1-C274A* cells are shown in blue on the left side; proteins up- regulated in nse1-C274A cells are shown in red on the right side; the position of specific protein is shown in yellow in the graph. **C**. Venn diagram comparing ubiquitin sites identified in this work, in Swaney et al ^1^ and in Tong et al ^2^. **D**. Gene ontology analysis of ubiquitinated sites significantly downregulated in *nse1-C274A* cells.

**Supplementary Figure 2.**
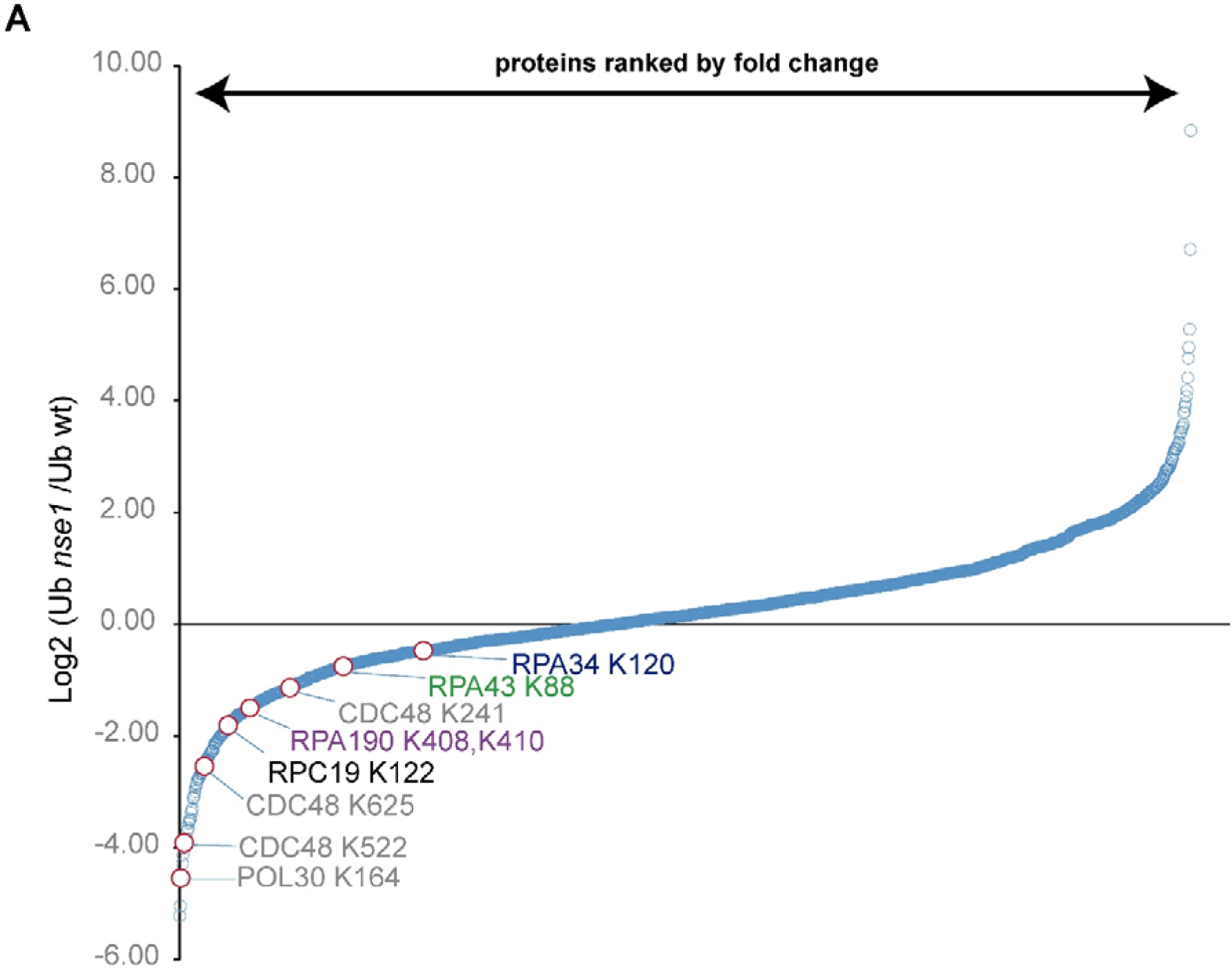
Ubiquitin sites in RNA polymerase I subunits. **A.** Waterfall plot showing the log2 fold change in ubiquitin site quantification from *nse1-C274A* (*nse1*) that significantly change relative to wild type (wt) nuclear extracts. Ubiquitin sites in RNAP I subunits are indicated; sites in Cdc48 and Pol30 are indicated as reference.

**Supplementary Figure 3.**
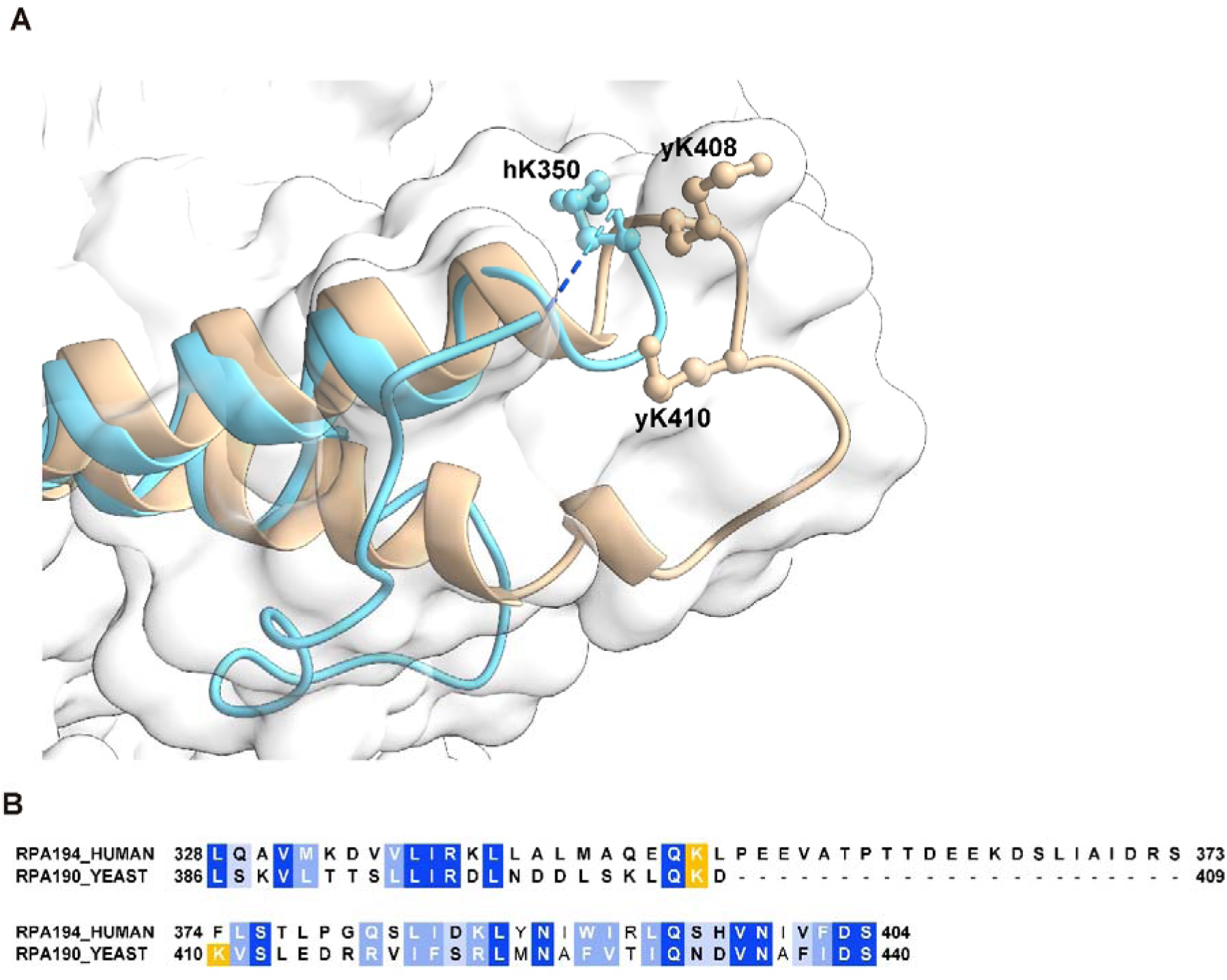
Structure and sequence alignment of yeast Rpa190 and human Rpa194. **A**. Structural alignment of clamp domains from yeast Rpa190 (in wheat; PDB 6H68 ^3^) and human Rpa194 (in cyan; PDB 7OB9 ^4^). The position of ubiquitinated residues, K408 and K410 in the yeast protein (yK408, yK410) and K350 in the human protein (hK350) are indicated. **B**. Sequence alignment of the clamp domain in the large subunit of the RNAP I complex. Dark blue indicates identical residues, light blue similar residues. Target lysines in the yeast protein (this study) and the human protein ^5^ are shown in yellow. .

**Supplementary Figure 4.**
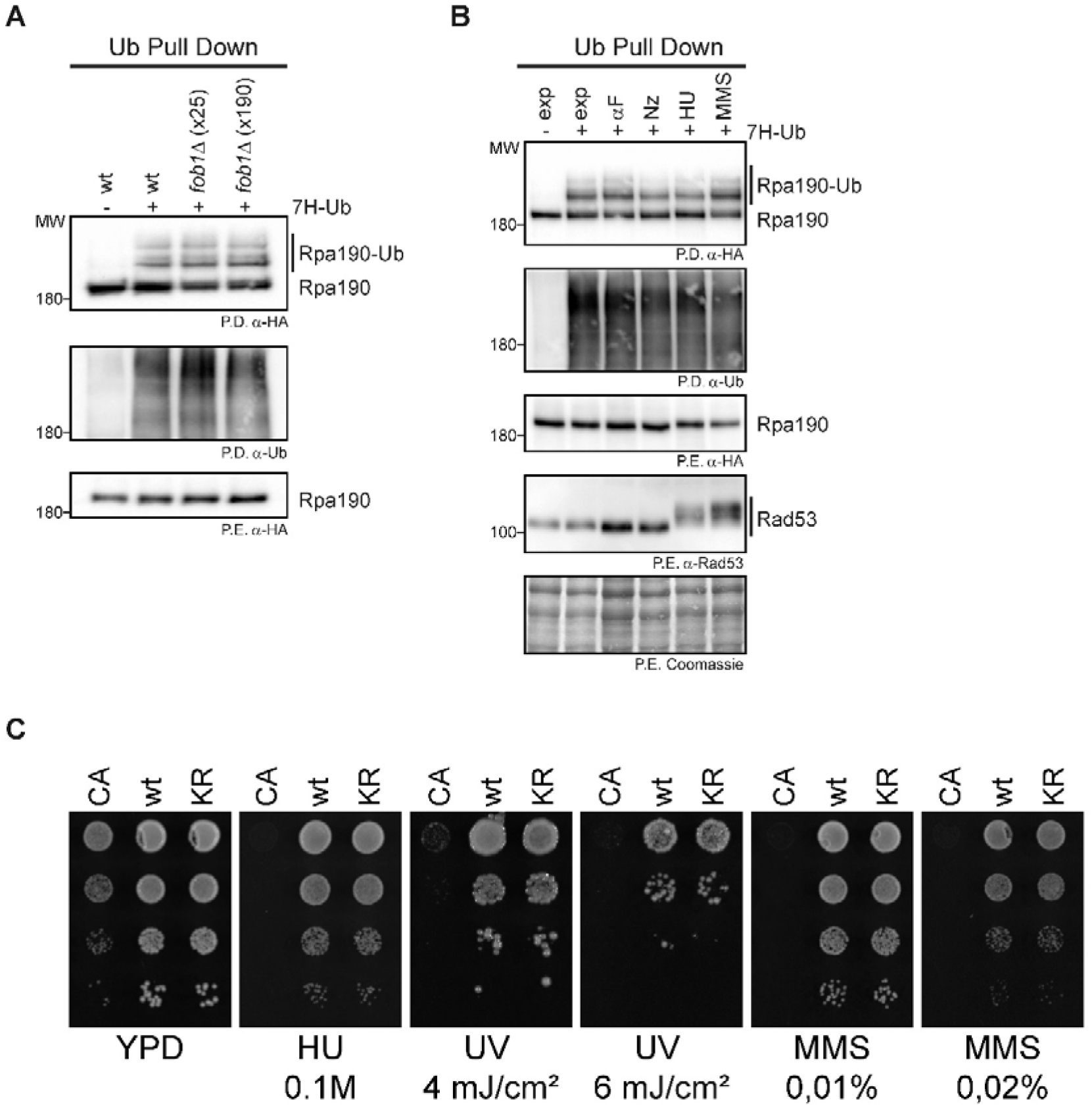
Rpa190 ubiquitination is not affected by rDNA copy number or cell cycle stage, and is not required for repair of UV-induced DNA damage. **A.** Exponentially growing cultures of wild type (wt), *fob1Δ* cells carrying 25 (x25) or 190 (x190) copies of the rDNA, a C-terminal 6xHA tag on *RPA190* at its endogenous location, and expressing (+) or not (-) a 7xHis tagged version of ubiquitin (7H-Ub) from the *CUP1* promoter, were collected; ubiquitinated species from protein extracts (P.E.) were pulled down (P.D.) under denaturing conditions with NiNTA beads. Rpa190-6HA species were detected with anti-HA antibodies. **B**. Exponentially (exp) growing cultures of wild type cells carrying a C-terminal 6xHA tag on *RPA190*, and expressing (+) or not (-) a 7xHis tagged version of ubiquitin (7H-Ub) from the *CUP1* promoter, were arrested in G1 with alpha factor (αF), in G2/M with nocodazole (Nz), in S phase with hydroxyurea 0.2 M (HU) or treated with MMS 0.02% to alkylate and damage DNA (MMS). Samples were processed as in A for ubiquitin pull down analysis. **C**. Growth test analysis wild type (wt) and *rpa190-KR* (KR) cells; 10-fold serial dilutions of the liquid cultures were spotted on YPD, supplemented or not with the indicated concentrations of hydroxuyrea (HU) or MMS or irradiated with the indicated doses of UV light. Plates were incubated at 30°C for 3 days. The *nse1-C274A* (CA) strain was included as a control for DNA damage sensitivity.

**Supplementary Figure 5.**
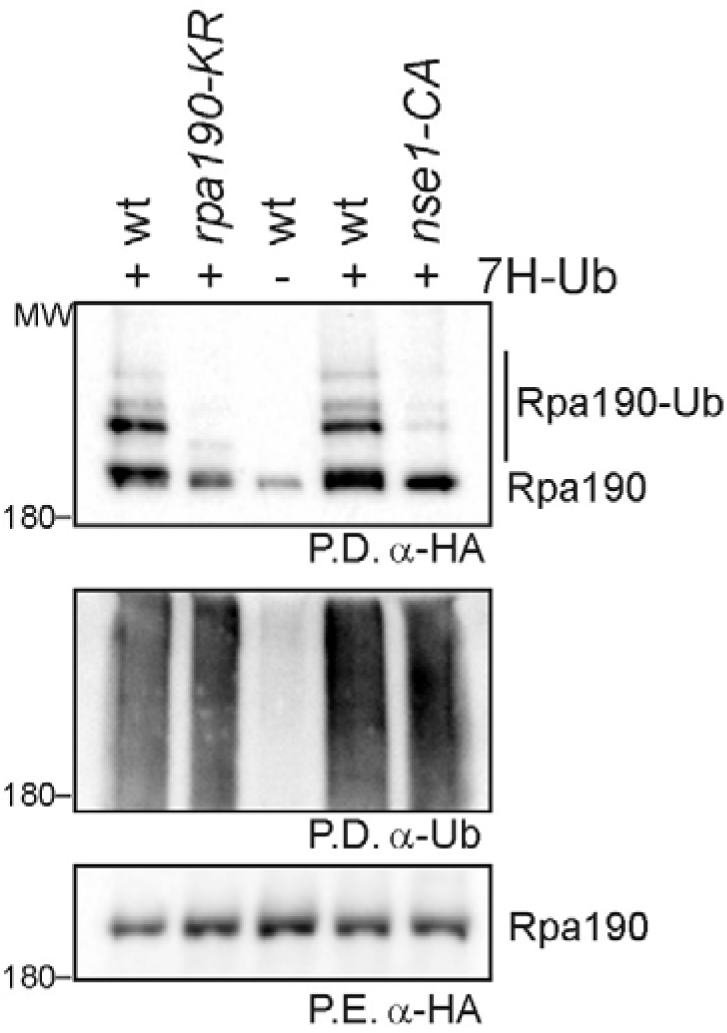
Rpa190 ubiquitination impairment is similar in *nse1-C274A* and *rpa190-KR* cells. Pull down analysis of Rpa190 ubiquitination in wild type (wt), *rpa190-KR* and *nse1-CA* cells; ubiquitin was pulled down (P.D.) under denaturing conditions from yeast protein extracts (P.E.) to purify ubiquitinated species.

**Supplemenetary Table 1.** Relevant genotype of yeast strains used in this study.

| NAME | GENOTYPE | PLASMIDS | FIGURE |
| --- | --- | --- | --- |
| EMY74.7 | *MATa his3-Δ1 leu2-3,112 trp1Δ ura3-52* |  | 1 |
| YRP787 | *MATa his3-Δ1 leu2-3,112 trp1Δ ura3-52 nse1::hisG pCEN-LEU2-nse1(C274A)* |  | 1 |
| YJC5738 | *MATa his3-Δ1 leu2-3,112 trp1Δ ura3-52 RPL24B-6HA:natNT2* | p2772, p2773 | 2 |
| YJC5740 | *MATa his3-Δ1 leu2-3,112 trp1Δ ura3-52 nse1::hisG pCEN-LEU2-nse1(C274A) RPL24B-6HA:natNT2* | p2772, p2773 | 2 |
| YJC5750 | *MATa his3-Δ1 leu2-3,112 trp1Δ ura3-52 ERG9-6HA:natNT2* | p2772, p2773 | 2 |
| YJC5831 | *MATa his3-Δ1 leu2-3,112 trp1Δ ura3-52 nse1::hisG pCEN-LEU2-nse1(C274A) ERG9-6HA:natNT2* | p2772, p2773 | 2 |
| YJC5746 | *MATa his3-Δ1 leu2-3,112 trp1Δ ura3-52 CDC73-6HA:natNT2* | p2772, p2773 | 2 |
| YJC5829 | *MATa his3-Δ1 leu2-3,112 trp1Δ ura3-52 nse1::hisG pCEN-LEU2-nse1(C274A) CDC73-6HA:natNT2* | p2772, p2773 | 2 |
| YGB5768 | *MATa his3-Δ1 leu2-3,112 trp1Δ ura3-52 YHB1-6HA:HIS3MX6* | p2772, p2773 | 2 |
| YGB5771 | *MATa his3-Δ1 leu2-3,112 trp1Δ ura3-52 nse1::hisG pCEN-LEU2-nse1(C274A) YHB1-6HA:HIS3MX6* | p2772, p2773 | 2 |
| YJC5838 | *MATa his3-Δ1 leu2-3,112 trp1Δ ura3-52 CDC48-6HA:natNT2* | p2772, p2773 | 2 |
| YJC5840 | *MATa his3-Δ1 leu2-3,112 trp1Δ ura3-52 nse1::hisG pCEN-LEU2-nse1(C274A) CDC48-6HA:natNT2* | p2772, p2773 | 2 |
| EMY74.7 | *MATa his3-Δ1 leu2-3,112 trp1Δ ura3-52* | p2772, p2773 | 2 |
| YRP787 | *MATa his3-Δ1 leu2-3,112 trp1Δ ura3-52 nse1::hisG pCEN-LEU2-nse1(C274A)* | p2772, p2773 | 2 |
| YNC2839 | *MATa his3Δ1 leu2Δ0 met15Δ0 ura3Δ0 RPA190-6HA:natNT2* | p2772, p2773 | 3B |
| YEI4864 | *MATa his3Δ1 leu2Δ0 met15Δ0 ura3Δ0 RPA190-K408R,K410R-6HA:natNT2* | p2772, p2773 | 3B |
| Y1838 | *MATa ade2-1 can1-100 his3-11,15 leu2-3,112 trp1-1 ura3-1 RAD5+* | pEI4637, pTR5024 | 3C |
| 61SC33.09 | *MATa ade2-1 can1-100 his3-11 leu2-3,112 trp1-1 ura3-1 RAD5+ bar1::LEU2 ubp10::hphMX4* | pEI4637, pTR5024 | 3C |
| Y1838 | *MATa ade2-1 can1-100 his3-11,15 leu2-3,112 trp1-1 ura3-1 RAD5+* | p2772, p2773, pEI4637, pTR5024 | 3D |
| 61SC33.09 | *MATa ade2-1 can1-100 his3-11 leu2-3,112 trp1-1 ura3-1 RAD5+ bar1::LEU2 ubp10::hphMX4* | p2773, pEI4637, pTR5024 | 3D |
| YNC2839 | *MATa his3Δ1 leu2Δ0 met15Δ0 ura3Δ0 RPA190-6HA:natNT2* | p2772, p2773 | 3E |
| YNC2839 | *MATa his3Δ1 leu2Δ0 met15Δ0 ura3Δ0 RPA190-6HA:natNT2* |  | 3F |
| YEI4864 | *MATa his3Δ1 leu2Δ0 met15Δ0 ura3Δ0 RPA190-K408R,K410R-6HA:natNT2* |  | 3F |
| Y1838 | *MATa ade2-1 can1-100 his3-11,15 leu2-3,112 trp1-1 ura3-1 RAD5+* | pEI4637, pTR5024 | 3G |
| 61SC33.09 | *MATa ade2-1 can1-100 his3-11 leu2-3,112 trp1-1 ura3-1 RAD5+ bar1::LEU2 ubp10::hphMX4* | pEI4637, pTR5024 | 3G |
| YEI3926 | *MATa his3-Δ1 leu2-3,112 trp1Δ ura3-52 RPA190-6HA:kanMX4* | p2772, p2773 | 4A |
| YEI4753 | *MATa his3-Δ1 leu2-3,112 trp1Δ ura3-52 nse1::hisG pCEN-LEU2-nse1(C274A) RPA190-6HA:natNT2* | p2773 | 4A |
| YNC2839 | MATa his3Δ1 leu2Δ0 met15Δ0 ura3Δ0 RPA190-6HA:natNT2 | p2772, p2773 | 4B |
| YEI4823 | nse5-ts3:kanMX4 MATa his3Δ1 leu2Δ0 ura3Δ0 met15Δ0 RPA190-6HA:natNT2 | p2773 | 4B |
| YGB5754 | nse1-16:kanMX4 MATa his3Δ1 leu2Δ0 ura3Δ0 met15Δ0 RPA190-6HA:HIS3MX6 | p2773 | 4B |
| YEI4664 | MATalpha can1Δ::MFA1pr-HIS3 lyp1Δ ura3Δ0leu2Δ0 his3Δ1 met15Δ0 RPA190-6HA:natNT2 | p2772, p2773 | 4C |
| YTR5618 | MATalpha can1Δ::MFA1pr-HIS3 lyp1Δ ura3Δ0leu2Δ0 his3Δ1 met15Δ0 RPA190-6HA:natNT2 rad6::kanMX4 | p2773 | 4C |
| YTR5620 | MATalpha can1Δ::MFA1pr-HIS3 lyp1Δ ura3Δ0leu2Δ0 his3Δ1 met15Δ0 RPA190-6HA:natNT2 mms2::kanMX4 | p2773 | 4C |
| YNC2839 | MATa his3Δ1 leu2Δ0 met15Δ0 ura3Δ0 RPA190-6HA:natNT2 | p2772, p2773 | 4D |
| YEI3959 | MATa his3Δ1 leu2Δ0 met15Δ0 ura3Δ0 RPA190-6HA:natNT2 mms21ΔC::hph | p2773 | 4D |
| YNC2839 | MATa his3Δ1 leu2Δ0 met15Δ0 ura3Δ0 RPA190-6HA:natNT2 |  | 4E |
| YSB2701 | MATa his3Δ1 leu2Δ0 met15Δ0 ura3Δ0 6HF-SMT3:kanMX6 RPA190-6HA:natNT2 |  | 4E |
| YTR5776 | MATalpha can1Δ::MFA1pr-HIS3 lyp1Δ ura3Δ0leu2Δ0 his3Δ1 met15Δ0 rpa190-K408R,K410R-6HA:natNT2 6HF-SMT3:KanMX6 |  | 4E |
| YSB2708 | MATa his3Δ1 leu2Δ0 met15Δ0 ura3Δ0 6HF-SMT3:kanMX6 mms21Dc::hphMX4 RPA190-6HA:natNT2 |  | 4E |
| YTR1437 | Mata bar1Δ leu2-3,112 his3-Δ200 trp1-Δ63 lys2-801 pep4 ade2-1::TetR-YFP:ADE2 ChrXII(487Kb)::tetO(5.6Kb):HIS3 ura3-52::ADH1-OsTIR1-9Myc:URA3 smc5-AID:kanMX4 |  | 5 |
| YTR5371 | Mata bar1Δ leu2-3,112 his3-Δ200 trp1-Δ63 lys2-801 pep4 ade2-1::TetR-YFP:ADE2 ChrXII(487Kb)::tetO(5.6Kb):HIS3 ura3-52::ADH1-OsTIR1-9Myc:URA3 smc5-AID:kanMX4 ubp10::hphMX4 |  | 5 |
| YTR5373 | Mata bar1Δ leu2-3,112 his3-Δ200 trp1-Δ63 lys2-801 pep4 ade2-1::TetR-YFP:ADE2 ChrXII(487Kb)::tetO(5.6Kb):HIS3 ura3-52::ADH1-OsTIR1-9Myc:URA3 smc5-AID:kanMX4 rpa190-K408R,K410R-6ha:hphNT1 |  | 5 |
| EMY74.7 | MATa his3-Δ1 leu2-3,112 trp1Δ ura3-52 |  | S1A |
| YRP787 | MATa his3-Δ1 leu2-3,112 trp1Δ ura3-52 nse1::hisG pCEN-LEU2-nse1(C274A) |  | S1A |
| YCC4680 | MATa leu2-3,112 ura3-1 his3-11,15 trp1-1 ade2-1 can1-100 RPA190-6HA:natNT2 |  | S4A |
| YEI3950 | MATa leu2-3,112 ura3-1 his3-11,15 trp1-1 ade2-1 can1-100 fob1::hphMX4 (rDNAx25) RPA190-6HA:kanMX |  | S4A |
| YEI3951 | MATa leu2-3,112 ura3-1 his3-11,15 trp1-1 ade2-1 can1-100 fob1::hphMX4 (rDNAx190) RPA190-6HA:KanMX |  | S4A |
| YNC2839 | MATa his3Δ1 leu2Δ0 met15Δ0 ura3Δ0 RPA190-6HA:natNT2 |  | S4B |
| YGB5752 | MATalpha can1Δ::MFA1pr-HIS3 lyp1Δ ura3Δ0 leu2Δ0 his3Δ1 met15Δ0 nse1-C274A:kanMX4 |  | S4C |
| Y2771 | MATalpha can1Δ::MFA1pr-HIS3 lyp1Δ ura3Δ0 leu2Δ0 his3Δ1 met15Δ0 |  | S4C |
| YEI5022 | MATalpha can1Δ::MFA1pr-HIS3 lyp1Δ ura3Δ0 leu2Δ0 his3Δ1 met15Δ0 RPA190-K408R K410R-6HA:natNT2 |  | S4C |
| YNC2839 | MATa his3Δ1 leu2Δ0 met15Δ0 ura3Δ0 RPA190-6HA:natNT2 | p2772, p2773 | S5 |
| YEI4864 | MATa his3Δ1 leu2Δ0 met15Δ0 ura3Δ0 RPA190-K408R,K410R-6HA:natNT2 | p2773 | S5 |
| YNC2839 | MATa his3Δ1 leu2Δ0 met15Δ0 ura3Δ0 RPA190-6HA:natNT2 | p2773 | S5 |
| YEI4864 | MATa his3Δ1 leu2Δ0 met15Δ0 ura3Δ0 RPA190-K408R,K410R-6HA:natNT2 | p2773 | S5 |

**Supplemenetary Table 2.**
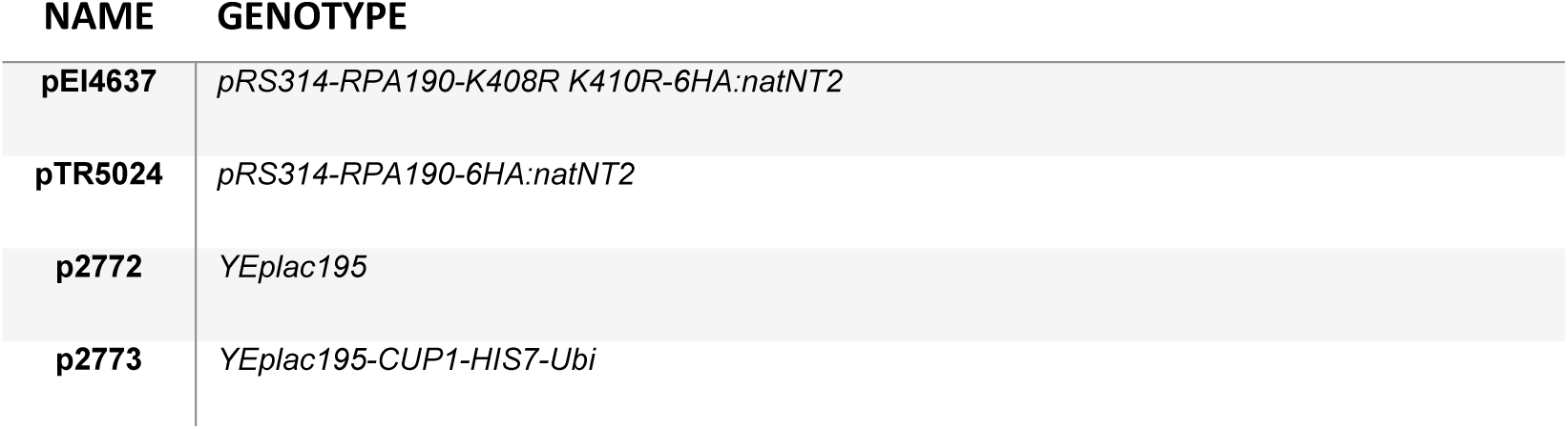
Relevant genotype of plasmids used in this study.

## Notes

### Competing Interest Statement

The authors have declared no competing interest.

